# Pathogen priming reveals host immune training and microbiome conditioning in corals

**DOI:** 10.64898/2026.04.07.716734

**Authors:** Matteo Monti, Neus Garcias-Bonet, Francisca C. García, Erika P. Santoro, Ghaida Aljuaid, Kevin Schuster, Chakkiath Paul Antony, Marco Casartelli, Laura Beenham, Aurora Giorgi, Luigi Colin, Christian R. Voolstra, Raquel S. Peixoto

## Abstract

In several species, vaccine-like approaches, where hosts are primed through controlled pathogen exposure, have proven effective in enhancing responses to subsequent infections. This principle remains unexplored in corals. Here, we demonstrate that chronic exposure to non-lethal concentrations of live or inactivated *Vibrio coralliilyticus* primes the coral holobiont to counter subsequent infections under heat stress. Non-primed corals experienced greater heat stress pathogen-driven bleaching and a significant decline in their photosynthetic efficiency compared to primed samples. These results were linked to microbiome conditioning and host gene expression modulation, including a layered, fine-tuned immune and cellular response to microbial invasion. This proof-of-concept challenges strictly innate immune responses in corals and positions immune priming and microbiome conditioning as integrated mechanisms of coral holobiont resilience. Together, these findings can contribute to redefine coral immunity concepts and lay the groundwork for developing new microbiome-based strategies to enhance coral health for reef conservation under climate change.

**LAY SUMMARY:** Coral reefs are increasingly threatened by rising seawater temperatures and disease. Unlike vertebrates, corals do not possess a classical adaptive immune system, so they have been thought to rely only on innate defenses. However, our study shows that corals may be more capable than previously believed. We demonstrate that exposing corals to non-lethal doses of a widespread bacterial pathogen can ‘train’ them to better withstand subsequent infections under heat stress.

Corals that were not pre-exposed suffered more pathogen caused bleaching and showed a stronger decline in physiological performance, while those that were primed were more resilient.

This improved resistance appears to come from two coordinated processes. First, the coral’s associated bacterial community shifted in a way that seems to help protect the host. Second, the coral itself adjusted its gene expression, mounting a more effective and controlled response to infection. These findings suggest that corals may be able to ‘remember’ past exposures and respond more effectively to future infections, even without a traditional adaptive immune system, providing a foundation for developing new strategies to support coral resilience.

## INTRODUCTION

Microbial communities are integral to the development, homeostasis, and resilience of multicellular organisms (1–3). Across ecosystems, host-associated microbiomes function not only as nutritional and immunological partners, but also as dynamic interfaces between hosts and their environments (4). Understanding how microbial communities assemble, stabilize, and respond to perturbations, such as pathogen invasion or environmental stress, is central to predicting host health and disease outcomes (5–7). A growing body of evidence suggests that host-microbiome interactions induce a form of ecological memory, whereby past exposures shape future responses through both microbial community shifts and host physiological conditioning (8–10). The concepts of microbiome ecological memory and adaptive microbiome flexibility (11–14) extend our understanding of microbial assemblages as active contributors to host defense (15–17) through their ability to be restructured by specific prior stimuli, moving beyond earlier views of them as static passengers. Microbiome reprogramming, by which specific exposures lead to lasting changes in microbial composition and function, is emerging as a foundational principle in host-microbiome biology, with implications for understanding holobiont evolution and adaptive responses in a changing world (13,18–20).

Corals exemplify the holobiont paradigm, relying on coordinated interactions with diverse symbiotic partners, including bacteria, archaea, viruses, fungi, and photosynthetic dinoflagellates, to maintain homeostasis and cope with environmental stress (21–23). Pathogenic infections can lead to the disruption of this finely tuned system leading to cascading effects such as the breakdown of the coral-algal symbiosis, coral tissue loss and eventually mortality.

Although the main global driver of coral bleaching and mortality is heat stress associated with marine heatwaves, in recent years coral diseases led to devastating outbreaks driving foundational reef species to collapse (e.g., the Caribbean reef-building elkhorn and staghorn corals, *Acropora palmata* and *Acropora cervicornis*, respectively) while contributing to the unprecedented decline of coral reef ecosystems worldwide (24–26).

Management strategies for controlling wildlife diseases, including those affecting corals, have traditionally relied on the use of antibiotics (27–30). However, their widespread and often inappropriate use, particularly when applied in complex and highly biodiverse systems such as coral reefs, may contribute to the emergence of antibiotic-resistant pathogens which undermines the efficacy of treatments across a growing spectrum of infections. This has, therefore, become a major concern not only for both human and wildlife health(3), but also for the stability of host-associated and environmental microbial communities that underpin reef ecosystem functioning (31–34). In response, inducing resistance against emerging pathogens has been demonstrated successfully through priming treatments in several animal taxa including mammals, fish, and amphibians, by conditioning the host for future infections (35–37).

As for all invertebrates, cnidarians (including corals) are thought to rely solely on innate immunity, involving pattern recognition receptors (PRRs) and cellular or humoral responses (38–40). However, growing evidence challenges this view suggesting that cnidarians immunity is much more complex than originally believed. For example, it has been demonstrated that the sea anemone *Exaiptasia pallida* has a complex innate immunity repertoire (41), harbors complex immunity responses to heat stress distinct from vertebrates (42), and when exposed to sub-lethal doses of a pathogen prior to lethal exposure showed higher survivorship compared to anemones encountering the infectious agent for the first time (43). The latter suggests that cnidarians may possess a protective priming defense mechanism against subsequent pathogen invasion. This responsive and adaptable interplay between host and its microbial partners opens new avenues for understanding disease resistance and increasing resilience in a rapidly changing ocean. Although our understanding of putative etiological agents, disease vectors and transmission mechanisms remains limited, the gram-negative bacterium *Vibrio coralliilyticus* has been identified as a widespread temperature-dependent coral pathogen (44,45). Specifically, *V. coralliilyticus* elicits disease symptoms across multiple scleractinian hosts including members of the genera *Acropora* and *Pocillopora*; its virulence is thermally modulated, with seawater temperatures above 26 – 27 °C inducing the expression of extracellular metalloproteases, toxins and secretions systems that damage the endosymbiotic dinoflagellates and coral tissue resulting in bleaching, leading ultimately to tissue loss and mortality (46–50). The pathogenicity of *V. coralliilyticus* has been determined predominantly only in a few strains such as BAA-450, P1, OCN008 and OCN014 (46,51,52), yet this bacterium is also isolated from apparently healthy colonies, consistent with an opportunistic lifestyle (53–55). These attributes, temperature-tunable virulence, reproducible disease phenotype and global occurrence, make *V. coralliilyticus* a model microorganism in coral disease research to investigate host-pathogen interactions, microbiome perturbation and host immune-like responses.

To assess the prospect of pathogen priming, we sought to use sublethal doses of *V. coralliilyticus*, either live or chemically inactivated, to prime fragments of the Red Sea reef-building coral *Pocillopora favosa* [formerly *P. verrucosa* (56)] by combining a coral mesocosm experiment with a multi-omics approach. This framework enabled us to evaluate whether prior exposure to the pathogen can reconfigure the coral associated bacterial communities and condition the host defenses, to ameliorate the stress response to subsequent pathogen exposure coupled with increasing temperatures.

## MATERIALS AND METHODS

### Sampling procedures

Four visually healthy coral colonies (i.e., no visible signs of stress, bleaching, disease or necrosis) of *Pocillopora favosa* were sampled on February 2024 by SCUBA diving from Al-Fahal reef (22°18’18.4"N; 38°57’52.5"E) in the central Red Sea (57), at a depth ranging between 6.00 and 10.00 meters, seawater temperature of 26.5 °C, and located at least 10 meters apart from each other to reduce the likelihood of obtaining clonal propagates. All colonies were transported in coolers filled with natural seawater until processing (≤ 3 h) at the Coastal and Marine Resources Core Lab (CMR) of the King Abdullah University of Science and Technology (KAUST, Thuwal, Saudi Arabia). All coral husbandry and experimentation were done under the supervision of KAUST’s Institutional Biosafety and Bioethics Committee (IBEC, Research protocol 22IBEC003).

At CMR, each *P. favosa* colony was fragmented in 36 nubbins (3-5 cm; N_tot_ = 144) using a diamond circular saw (Gryphon Corp., USA), using standard procedures (58) and left to heal and acclimate for three days in a 300-liters open-system healing aquarium receiving natural seawater and equipped with four EcoTech Marine Radion XR15 full spectrum led lights (EcoTech Marine, USA) set to a maximum intensity of 200 µmol of photons m^-2^s^-1^ between 12:00 and 1:00 pm, reflecting the lower bound of *in situ* midday conditions at a depth of 6 to 10 meters in Al Fahal reef, and a 12-hour day/night cycle changing at 7:00 am and 7:00 pm. During the three-days acclimation period the seawater temperature was gradually reduced from 26.5 to 25.5 °C (0.5 °C decrease in the first two days), while the water parameters were monitored daily for all the experiment timeframe with a YSI ProDSS multiparameter digital water quality meter (YSI Incorporated, USA) and maintained stable at pH 7.9 – 8.1, salinity 40.00 – 40.20 PSU, and DO 6.15 – 6.21 mg/L.

### Mesocosm experimental design, coral priming, and exposure to *V. coralliilyticus* infection at increasing temperature

The experimental mesocosm used for this priming experiment consisted of three independent 300-liters water baths, in which four individual 10-liters glass cubic experimental aquariums from each treatment were placed. Each completely individualized aquarium was an independent true biological replica, equipped with a protein skimmer to remove excess nutrients in the water and a circulation pump to ensure constant water movement, kept at a stable temperature of 25.5 °C, 12-hours photoperiod and water parameters maintained as in the previously described healing aquarium. Water changes (50 % of the total volume for each tank) occurred every day in the morning 2 hours after the start of the daily photoperiod using natural seawater from the healing tank. The water in the water baths was homogenized by two circulation pumps to maintain a homogeneous temperature of 25.5 °C, and there was no water exchange between the experimental aquariums and the water baths. Each water bath was equipped with a thermostat that monitored and regulated temperature by activating heaters as needed. Heating was provided by two 100-W heaters per bath.

After healing and acclimation, 12 coral fragments (three from each coral colony) were placed in each of the 10-liters experimental aquariums and left to acclimate for three days. Immediately after the three-days acclimation period (T0), one fragment per colony from each of the 12 treatment aquariums (N_tot T0_ = 48) were snap-frozen in liquid nitrogen and stored at -80 °C to be processed and used as a baseline for microbiome and host gene expression analysis.

Four treatments were compared to test the potential and efficacy of coral prophylaxis: 1) C: control corals, samples not subjected to priming nor exposed to subsequent pathogen infection at high temperature; 2) NP: samples not subjected to priming but exposed to subsequent pathogen infection at high temperature; 3) LP: priming employing live pathogen (10^4^ live cells/ml of *V. coralliilyticus* every three days after T0) and subsequent pathogen infection at high temperature; 4) DP: priming employing chemically inactivated pathogen (10^4^ chemically inactivated cells/ml of *V. coralliilyticus* every three days after T0) and subsequent pathogen infection at high temperature. *Vibrio coralliilyticus* strain BAA-450 (referred as VBAA450) was chosen as target pathogen for prophylaxis and subsequent infection because of its demonstrated ability to cause bleaching and tissue loss in other species of the genus *Pocillopora* [i.e., *P. damicornis* (51)].

To obtain live pathogen cells for coral priming, fresh VBAA450 cultures were prepared for each priming inoculation (every two days) in sterile marine broth (SMB, Marine Broth 2216, Himedia Laboratories) and incubated at 30 °C/120 rpm shaking for 24 hrs. Cell concentration for each culture was estimated using a Multiskan SkyHigh Microplate Spectrophotometer (Thermo Scientific™, USA) to measure optical density at 600 nm (OD_600_) corresponding to approximately 10^9^ cells/ml. Each VBAA450 culture was then spun down to pellet cells, supernatant was removed, and cells were washed three times and re-suspended in sterile 3.5 % saline solution (3.5 % SSS). Finally, a serial dilution step to 10^7^ VBAA450 cells/ml cultures was performed to achieve a final priming inoculation concentration of 10^4^ pathogen cells/ml in the 10-liters treatment aquaria. Similarly, to obtain chemically inactivated pathogen cells for the second priming treatment, after culturing and supernatant removal, VBAA450 cells were resuspended in a 1.0 % formalin solution and left at 4 °C overnight. After the overnight incubation at 4 °C, chemically inactivated VBAA450 cultures were washed three times and re-suspended in sterile 3.5% SSS to an OD_600_ of 10^7^ cells/ml. The complete inactivation of VBAA450 cells was confirmed by plating 100 µl of the resuspended cell culture onto triplicate sterile Petri dishes containing 20 ml of marine agar (MA, Difco^TM^ Marine Agar 2216), incubated for 24 hrs at 30 °C, on which no bacterial colony forming units grew.

Priming regimen was established for 21 days by inoculating 10 ml of live and chemically inactivated 10^7^ VBAA450 cells/ml into the 10 liters DP and LP treatment aquaria every three days (final concentration: 10^4^ VBAA450 cells/ml). C and NP aquaria were treated in a similar manner by inoculating 10 ml of placebo 3.5 % SSS. Protein skimmers were turned off for three hours from inoculation and 50 % water changes were performed every other day for the whole priming timeframe. At the end of the 21-days priming phase (T1) one coral fragment per colony from each of the 12 treatment aquariums (N_tot T1_ = 48) were snap-frozen in liquid nitrogen and stored at -80 °C to be processed.

Following the VBAA450 priming treatments (T1), a 100 % water change was performed in each of the 12 treatment tanks and left for a resting period of four days in which 50 % water changes occurred daily. Subsequently, temperature in the three water baths containing the treatment aquaria was increased from 25.5 °C to 32.5 °C over the course of 10 days (0.7 °C increase/day) to trigger the temperature-dependent pathogen virulence. The target temperature of 32.5 °C was selected based on previous studies showing that this temperature, although elevated, does not elicit visible signs of bleaching in *P. verrucosa* when applied alone (59,60). While increasing temperature, coral samples in all the NP, LP and DP aquaria were exposed, every three days for 13 days, to the infection of live VBAA450 at a final concentration of 10^6^ cells/ml (fresh cultures of pathogen were obtained as described in the previous section). C aquaria did not receive any pathogen inoculation as they served as temperature controls to verify that potential coral bleaching was not caused by the temperature increase. After 15 days of infection regimen (T2) all the remaining coral fragments (N_tot T2_ = 48) were snap-frozen in liquid nitrogen and stored at - 80 °C to be processed. Both priming 10^4^ cells/ml and infection 10^6^ cells/ml final concentrations of VBAA450 were chosen based on the *Vibrio* inoculum size applied in several other infection experiments in corals and other invertebrates present in the scientific literature (51,52,61,62).

### Assessment of coral health

Coral health was assessed during the experiment using different proxies, including algal photosynthetic parameters as well as visual monitoring of stress signs, including chronic polyp retraction, paling, bleaching and tissue loss. We assessed the photochemical efficiency of the coral associated Symbiodiniaceae at T0, T1, at days 4 and 8 of VBAA450 infection (respectively at 28.3 °C and 31.1 °C), and at T2 by employing pulse-amplitude-modulated (PAM) fluorometry, which was used as a proxy for the health of the coral holobiont (63). To avoid interference from diurnal photoinhibition artifacts we performed measurements using a submersible diving-PAM system (Walz GmbH, Germany) one hour after sunset (20:00 h) by placing the 8-mm glass fiber optic probe above the oral disk of the coral polyps. All measurements were taken on the side and middle of the fragments to avoid measurements of the tips or base. The maximum quantum yield of PSII photochemistry was determined as *F_v_*/*F_m_*, where *F_v_* was obtained as *F_m_ –F_o_* (*F_o_* = maximum level of fluorescence detected under the modulated measuring light of the PAM [weak pulsed light <1 µmol photons m^−2^ s^−1^]; *F_m_* = maximum fluorescence level detected using a short saturating pulse of actinic light). The diving PAM was configured as follows: measuring light curve intensity = 6, measuring light frequency = 3, signal gain = 2, signal damping = 4, electron-transfer-rate-factor (ETR-Factor) = 0.85, and Actinic Light Factor = 1.

As *F_v_*/*F_m_* data resulted not normally distributed (Shapiro-Wilk: p < 0.05), changes in photosynthetic efficiency across treatments were determined using a general linear mixed-effects model (GLMM) with a ‘Gamma’ family distribution with a ‘log link’ function using the ‘glmer’ function in the lme4 package in R (64) per each time *F_v_*/*F_m_* data were collected. Coral colonies and tank replicates were accounted for as random effects on the intercept. *F_v_*/*F_m_* measurements were included in the model as response variable while ’treatment’ with four levels (C, NP, LP, DP) represented the predictor variable. Overall changes in *F_v_*/*F_m_* across treatment levels at T0, T1, at days 4, 8 and 13 (T2) of VBAA450 infection were determined by ANOVA tests comparing GLMMs with their respective null models, followed by pairwise comparisons between treatments by applying the ‘emmeans’ R function on the fitted models using Tukey adjustment.

We also obtained photographs of the coral fragments which were taken at T1 (25.5 °C), at day 4 of VBAA450 infection (28.3 °C) and at T2 (32.5 °C) using an Olympus TG-6 camera to evaluate holobiont stress conditions. To quantify the brightness intensity, in Adobe Photoshop CC 2024, each coral fragment was digitally isolated from the background and converted to 8-bit grayscale using Photoshop’s default grayscale profile. For each fragment, the mean gray value (ranging from 0 = black to 255 = white) was automatically computed representing the unweighted average brightness across all pixels within the coral fragment [similarly to (65,66)]. These values were then used as an estimate to evaluate potential significant differences in pathogen induced bleaching severity between samples across treatments by employing GLMMs and pairwise comparisons between treatments as per *F_v_*/*F_m_* data.

### Coral microbiome assemblage

Coral associated microbiome was assessed by 16S rRNA gene amplicon sequencing analysis focusing exclusively on the composition and dynamics of bacterial taxa. Accordingly, for the purpose of this study we use the term microbiome to indicate host-associated bacterial community, unless otherwise specified. At each sampling time (T0, T1, and T2) we removed one entire fragment for each *P. favosa* colony from each of the 12 treatment aquaria (N_tot x sampling time_ = 48), which were then snap frozen in liquid nitrogen and placed at - 80 °C until processing. Approximately 0.30 g of each sample was incubated overnight with 700 µl of DESS buffer (20% dimethyl sulfoxide, 0.25 M ethylenediaminetetraacetic acid, and saturated sodium chloride (NaCl), with adjusted pH 8.0) before DNA extraction. Total DNA was extracted using the DNeasy^®^ Blood and Tissue kit (Qiagen, Germany) following the manufacturer’s instructions with a Gram-positive pre-treatment step, consisting of a 37 °C incubation for 30 minutes in enzymatic lysis buffer, and a 56 °C 3-4 h incubation in the kit lysis solution shaking at 650 rpm in a Thermomixer (ThermoFisher^®^, USA). The extracted DNA was quantified using a high sensitivity Qubit™ dsDNA assay kit (Invitrogen™, USA) and then the hypervariable V3 and V4 regions of the bacterial 16S rRNA gene were amplified in triplicate PCRs using the universal primers 341F 5’ CCTACGGGNGGCWGCAG 3’ and 785R 5’ GACTACHVGGGTATCTAATCC 3’. PCR reactions consisted of 12.50 µl of Kapa HiFi Hotstart Ready Mix (Roche Holding AG, Switzerland), 1.0 µl from primer F and R, 8 µl of PCR grade water and 2.5 µl of sample DNA standardized to a concentration of 5 ng/µl. Thermal cycling conditions were 95 °C for 3 min, followed by 35 cycles of 95 °C for 30 s, 62 °C for 30 s and 72 °C for 30 s, with a final 5 min extension at 72 °C. For each PCR replicate, negative reagent controls without template DNA were run. PCR products were verified and quantified by mixing their equal volume with 1X loading buffer (contained SYBR Green) and performing electrophoresis on 2 % agarose gels. PCR plates were then stored at - 20 °C until library preparation. PCR products were purified using a Qiagen Gel Extraction Kit (Qiagen, Germany). Sequencing libraries were generated with a NEBNext® Ultra™ II DNA Library Prep Kit (Cat No. E7645, New England Biolabs). The library quality was evaluated on a Qubit™ 2.0 Fluorometer (Thermo Scientific™) and Agilent Bioanalyzer 2100 system. Libraries were sequenced on a NovaSeq platform (Illumina), and 250 bp paired-end reads were generated.

The DADA2 pipeline (67) was used to process the 16S rRNA gene-based amplicon libraries. Briefly, the raw reads were decontaminated by phiX and adapter-trimmed using the “BBDuk” tool from the BBMap suite (68). PCR primers were then removed from the reads using the “cutadapt” tool (69) and the maxEE (maximum expected error) parameter at the “FilterAndTrim” step of DADA2 was set to 6 for the forward and reverse reads. After performing concatenation of the forward and reverse reads via the “justConcatenate” option in the “mergePairs” function of DADA2, the sequences were analyzed under the pseudo-pooling mode by following the standard DADA2 (version 1.22) workflow.

All analyses were performed in R using the functions in Phyloseq version 1.42.0 (70). Taxonomy was assigned to ASVs at 99% sequence identity using the SILVA-138 classifier (71). Reads identified as mitochondria, chloroplast, archaea, eukaryotes, and singletons were removed before the ASV table was rarefied to 100,000 reads (read depth determined by a ASVs accumulation curve; Supplementary Fig.1) using the ‘rarecurve’ function and the ‘rarefy_even_depth’ function in the Vegan package to avoid sequencing coverage bias (72).

**Figure 1.**
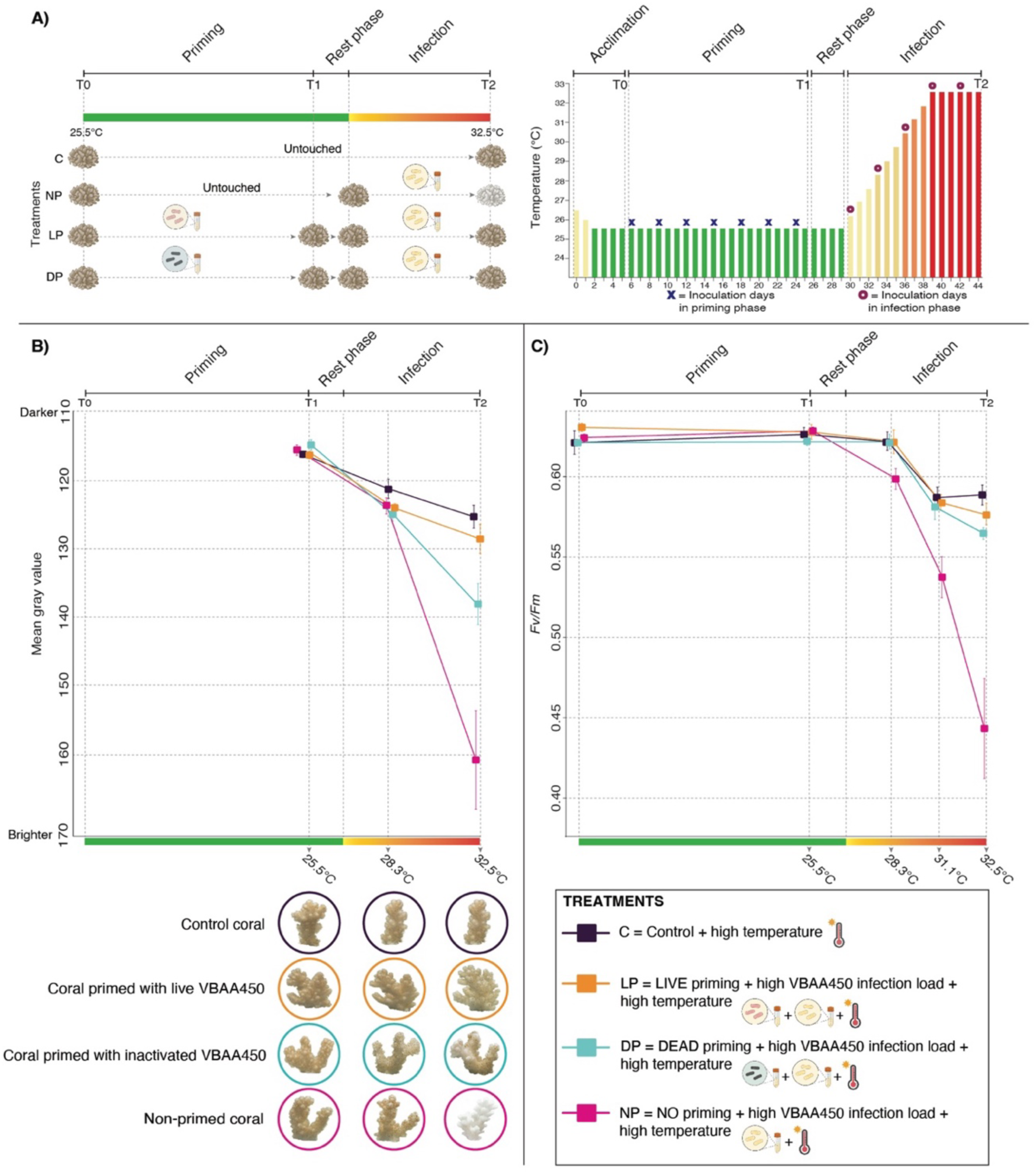
Coral priming experiment with *V. coralliilyticus* and coral physiological response to subsequent infection. A) Experimental design overview. B) Mean gray value (y axis) recorded as a proxy for pathogen-induced bleaching and photos of representative coral fragments taken at 25.5, 28.3 and 32.5 °C. The same coral fragments were followed over time within each treatment; however, photographic orientation was not always consistent across timepoints due to routine handling during *F_v_/F_m_* measurements and husbandry. C) Mean photosynthetic efficiency (*F_v_*/*F_m_* ratios; y axis) from coral fragments. Square symbols represent the mean value for each treatment (C = control samples; NP = non-primed samples; LP = live VBAA450 primed samples; DP = inactivated VBAA450 primed samples), and the error bars represent standard errors.

Alpha diversity was calculated using the ‘estimate_richness’ function in Phyloseq, with observed number of ASVs, Shannon H’, and Chao1 diversity indices. Statistical comparisons between treatments for alpha diversity metrics were calculated by implementing the Dunn test for multiple comparisons (previous testing for normal distribution of the data; Shapiro-Wilks, p-values < 0.05). The unweighted and weighted UniFrac distance matrices were used as metrics for beta diversity. The ‘betadisper’ command in the Vegan package was used to check for differences in homogeneity of variances. Principal coordinates analyses (PCoAs) were used to visualize patterns in beta diversity and permutational multivariate analysis of variance (PERMANOVA) was performed using the ‘adonis2’ function from the vegan R package on both unweighted and weighted Unifrac distance matrices to assess how treatment and timepoint affect the coral bacterial communities. To account for the non-independence of samples originating from the same colony, permutations were restricted within colony using a blocking (strata) design. The PERMANOVA model included treatment and timepoint as main effects, and significance was assessed using 99,999 permutations. Pairwise PERMANOVA tests were conducted using the ‘pairwise.adonis2’ function (73) to evaluate differences between specific treatment and timepoint combinations.

To confirm the presence of the VBAA450 in the coral microbiome after priming, the 16S rRNA gene sequence of *V. coralliilyticus* strain YB1 [= ATCC BAA450; (51,61)] was retrieved from there NCBI-nt database (NCBI Reference Sequence: NR_028014.1) and was used to query ASVs from each of our experimental groups at the different timepoints of the study at 100 % sequence identity. Finally, to identify differentially abundant ASVs across treatments at T1 and T2, the Analysis of the Composition of Microbiomes with Bias Correction 2 “ANCOM-BC2” (74) was implemented on total ASV counts (after removing reads identified as mitochondria, chloroplast, archaea, eukaryotes and singletons), using the Benjamini-Hochberg (BH) method to correct for false positives and an alpha of 0.05 for significance. An ASV was considered enriched or decreased significantly at a p_adj._ < 0.05 between treatments at T1 and T2.

### Coral host RNA extraction, sequencing, and differential gene expression analysis

Total RNA was extracted from approximately 0.20 - 0.30 g of randomly picked nubbins from four coral colonies (N_C_ = 4, N_NP_ = 4, N_LP_ = 4, N_DP_ = 4 at T0, T1 and T2; N_tot_ = 48) so that identical genotypes with the same life history were used for each experimental treatment. Frozen coral samples were placed in sterile 2.0 ml cryovials with ceramic beads filled with 1.0 ml of -20 °C cold urea-lithium chloride solution (8M urea, 3M LiCl) and incubated overnight shaking at 350 rpm at 4 °C. Coral tissue for each sample was then homogenized two times in a Qiagen TissueLyser II Bead Mill at 30Hz x 30 seconds and then the homogenate was transferred into a QIAshredder column (Qiagen, Germany) and centrifuged at 5,000 x g for 30 minutes at 4 °C. Following centrifugation, supernatant was discarded while tissue pellets were resuspended and incubated in 1.0 ml of TRIzol for 5 minutes at room temperature, subsequently hand-mixed, and incubated for 10 additional minutes with the addition of 200 µl of chloroform at room temperature. Samples were then centrifuged for 15 minutes at 12,000 x g / 4 °C to separate the TRIzol-chloroform solution into a lower ren phenol-chloroform phase, an interphase and a colorless upper aqueous phase containing the RNA. This aqueous phase was then transferred in a new 2.0 ml sterile tube with the addition of 1 volume of 70 % ethanol. Finally, RNA was extracted and eluted using the RNeasy mini kit (Qiagen, Germany) following manufacturer’s instructions. For each sample, RNA was quantified using a high sensitivity Qubit™ RNA assay kit (Invitrogen™, USA), checked for integrity and purity using an Agilent Bioanalyzer 2100 system and via a Nanodrop ND2000 spectrophotometer (Thermo Fisher™, USA).

Subsequently, 20 µl of each RNA sample was placed in a GenTegra-RNA 96-well microplate (NBS Scientific, USA) and left overnight to dry at room temperature before being sent for mRNA sequencing at BMK (Biomarker Technologies GmbH, DE). Sequencing was performed as 2x150 bp paired end on a Novaseq platform (Illumina).

mRNA sequencing of the 48 *P. favosa* tissue samples yielded a total of 1,071.14 M paired end reads (319.58 Gb), with an average yield of 6.66 Gb per sample, ranging from 20.5 to 87.2 million read pairs per sample after adapter trimming and quality control.

Approximately 72.84 % of reads aligned to the *P. verrucosa* ASM3666991v2 assembly genome [https://www.ncbi.nlm.nih.gov/datasets/genome/GCF_036669915.1/; (75)]. From a total of 32,915 genes, 3,820 represented non-functional pseudogenes, 569 tRNA genes, while 28,526 were assigned to annotated ones. Genome files, including the genomic FASTA and annotation files, were downloaded using the NCBI Datasets command-line tool (76). For compatibility with downstream RNA-Seq analysis pipelines, the downloaded GFF3 annotation file was converted to GTF format using AGAT (77). For functional annotation and downstream analyses, a custom organism annotation package was generated using the R package AnnotationForge (78). NCBI gene information files for *P. verrucosa* (taxon ID: 203993) were downloaded and used as input. The resulting annotation package (org.Pverrucosa.eg.db) was installed locally for use in Gene Ontology (GO) analyses. RNA data quality control, read trimming, alignment to the reference genome with a splice-aware aligner, and quantification of gene expression levels with STAR (79) and Salmon (80) was done using nf-core/rnaseq v3.18.0 (81) of the nf-core collection of workflows (82). Nf-core collection utilizes reproducible software environments from the Bioconda and Biocontainers projects (83,84). The pipeline was executed with Nextflow v24.10.5 (85). All downstream analyses were performed in R. To visualize global patterns in gene expression and sample relationships, ordination analyses were performed using principal component analysis (PCA). Raw gene count data from nf-core/rnaseq were imported and preprocessed with base R. The count data were log10-transformed to stabilize variance and reduce the influence of highly expressed genes. Ordination plots were generated for all samples and stratified by treatment and timepoint. In addition, principal component analysis (PCA) was performed on variance-stabilized transformed (VST) counts corresponding to all possible experimental pairwise comparisons. Differential expression analysis was performed using DESeq2 [v1.46.0; (86)] with the following parameters: lfcThreshold = 0, alpha = 0.05, cooksCutoff = FALSE, and independentFiltering = FALSE. The experimental treatments and sampling times were incorporated as explanatory factors in different DESeq2 models to capture distinct sources of transcriptomical variation, which was assessed using contrasts generated under different frameworks: (i) comparisons between treatments within the same timepoint and (ii) within-treatment longitudinal comparisons across timepoints to identify genes whose expression changed relative to baseline over the duration of the experiment. For each comparison, results were filtered for adjusted p-values (q-value) ≤ 0.05. Default p-value adjustment with the Benjamini-Hochberg false discovery rate (FDR) method was used. The number of differentially expressed genes, as well as the number of up- and down-regulated genes, was summarized for all contrasts defined within these analytical frameworks using custom functions. Annotated count tables were generated by merging DESeq2 results with gene annotation information. Gene Ontology (GO) enrichment analysis was performed with the package clusterProfiler [v4.14.6; (87)] to identify biological processes overrepresented among differentially expressed genes. Gene-to-GO term mappings were obtained from a custom annotation database (i.e., org.Pverrucosa.eg.db) generated for the *Pocillopora verrucosa* reference genome. Significantly differentially expressed genes (q-value < 0.05) with a minimum of fold change of 2 (absolute log2FC > 1) were used as input for GO enrichment analysis with the enrichGO function from the clusterProfiler package (88), focusing on Biological Process (BP) terms, and using the FDR method for multiple testing correction, with a q-value cutoff of 0.05. REVIGO (89) was used in R to cluster semantically redundant GO terms at a threshold of 0.3 for each treatment across timepoints using the package rrvgo (90) and treemaps per each treatment were used to represent REVIGO non redundant GO terms using the function ‘treemapPlot’ in the treemap package (91). To assess how treatment and timepoint affect gene expression, PERMANOVA was performed using the ‘adonis2’ function from the vegan R package. Euclidean distance matrices were computed from the transformed expression values for all samples. To account for the non-independence of samples originating from the same colony, permutations were restricted within colony using a blocking (strata) design. The PERMANOVA model included treatment and timepoint as main effects, and significance was assessed using 99,999 permutations. Pairwise PERMANOVA tests were conducted using the ‘pairwise.adonis2’ function to evaluate differences between specific treatment and timepoint combinations.

## RESULTS

### Priming treatments mitigate pathogen stress at high temperatures without compromising host health

As a first assessment, we performed visual inspection of each of the 144 coral fragments at T0 which revealed no sign of stress (e.g., changes in color, tissue loss or excessive mucus production, intended as the presence of visible mucus sloughing or sustained mucus release from the coral surface) suggesting full acclimation of all samples to the experimental tank conditions. No visual differences were also detected throughout the 21 days priming timeframe at 25.5 °C for both control (C) and non-primed (NP) samples, while we observed a transient retraction of coral polyps, characterized by the withdrawal of the tentacles into the calyx, that lasted approximately one hour after each inoculation of live and inactivated *Vibrio coralliilyticus* BAA-450 (from here on VBAA450) in the respective treatments (LP = live VBAA450 priming and DP = chemical inactivated VBAA450 priming). More in detail, analysis of the coral fragments’ photographs revealed no significant differences after priming at T1 in the mean gray value of each sample in pairwise comparisons between treatments and controls (Wald Z-test 25.5 °C, p_adj_ > 0.05; Fig. 1B; Supplementary Tables 1, 2). Moreover, photosynthetic efficiency (*F_v_*/*F_m_* values), indicative of the host-associated Symbiodiniaceae photosystem function did not differ across controls and treatments in any pairwise contrasts at both T0 and T1, with values remaining stable in both controls and treatments (Fig. 1C; Supplementary Tables 3, 4).

Nevertheless, during the post-priming VBAA450 infection phase, we recorded an overall significant effect of the different treatments on *P. favosa* brightness under combined pathogen exposure and thermal stress. Treatment effects emerged starting at 28.3 °C and intensified at higher temperatures (32.5 °C = T2). Generalized linear mixed models revealed a significant treatment effect at 28.3 °C (AIC = 270.06, χ² = 7.96, df = 3, *p* = 0.047) as well as 32.5 °C (AIC = 378.14, χ² = 48.11, df = 3, p < 0.001; Fig. 1B; Supplementary Tables 1, 2). Although we did not detect significant pairwise contrasts across treatments at 28.3 °C, at 32.5 °C coral fragments exposed to pathogen loads without previous priming (NP) showed significantly brighter coloration than coral fragments that were primed before exposure to pathogen (LP and DP) and those fragments that never encountered the pathogen for the whole experiment (C) [average coral fragments brightness: NP = 160.67 (±7.18); C = 125.28 (±1.67); DP = 137.98 (±3.00); LP = 128.51 (±2.12); Wald Z-tests, p_adj_ < 0.001]. Moreover, in contrast to the significant difference observed between the C and DP groups, no statistically significant difference in coral paling was detected between control corals and VBAA450 live-primed corals. Similarly, GLMMs on PAM data revealed overall significant differences in *F_v_*/*F_m_* values across treatments starting from 28.3 °C (AIC = -224.08, χ² = 11.78, df = 3, p = 0.082; Fig. 1C; Supplementary Tables 3, 4) with significant pairwise differences between NP treatment compared to C, DP and LP groups (Wald Z-tests, p_adj_ = 0.013, 0.143 and 0.015 respectively). Although a reduction of *F_v_*/*F_m_* values was recorded in all treatments, at 32.5 °C (T2) the non-primed group was the only one showing a significant drop in photosynthetic efficiency compared to the control (C) as well as to both priming treatments DP and LP (GLMM_T2_: AIC = -7.10, χ² = 9.47, df = 3, p = 0.024; pairwise contrasts: p_adj_ = 0.019, 0.040 and 0.0315 respectively). Notably, no tissue loss nor mortality were observed in any of the coral fragments at T2.

### Priming treatments alter the host-associated microbiome

Amplicon 16S rRNA gene sequencing revealed a significant restructuring effect following the different treatments across the timeframe of the experiment on the host-associated bacterial communities (hereafter referred to as the microbiome) both in terms of the presence/absence of ASVs (unweighted UniFrac Adonis _(treatment*sampling_time)_, R2 = 0.448, df = 6, F = 1.157, Pr (>F) = 0.044; Permdisp, Pr (>F) = 0.199), as well as in terms of their relative abundance (weighted UniFrac Adonis _(treatment*sampling_time)_, R2 = 0.034, df = 6, F = 2.140, Pr (>F) = 0.018; Permdisp, Pr (>F) = 0.196) (Fig. 2A, Supplementary Table 5). Temporal microbiome shifts shared across treatments reflect background acclimation effects, while priming effects were inferred only from between-treatment differences. As expected, at T0 (before priming) the host-associated bacterial assemblages did not show any pairwise significant differences between treatments, with alpha-diversity metrics further suggesting similarity of microbiomes (Dunn’s tests on ASVs number, Shannon and Chao1 diversity indices across treatments p > 0.05; Supplementary Tables 6, 7). Conversely, both priming treatments triggered a significant assemblage shift of the microbiomes associated with *P. favosa* samples both in terms of ASVs presence/absence as well as in terms of their relative abundance. Specifically, when performing pairwise comparisons at T1, beta diversity metrics showed significant differences between C and LP samples (unweighted UniFrac Adonis, df = 1, F = 1.591, R2 = 0.081, p_adj_ = 0.002; weighted UniFrac Adonis, df = 1, F = 5.933, R2 = 0.248, p_adj_ = 0.015), C and DP samples (unweighted UniFrac Adonis, df = 1, F = 1.917, R2 = 0.083, p_adj_ = 0.002; weighted UniFrac Adonis, df = 1, F = 4.690, R2 = 0.182, p_adj_ = 0.012), NP and LP samples (unweighted UniFrac Adonis, df = 1, F = 1.685, R2 = 0.090, p_adj_ = 0.004; weighted UniFrac Adonis, df = 1, F = 5.480, R2 = 0.244, p_adj_ = 0.008), and finally NP and DP coral fragments (unweighted UniFrac Adonis, df = 1, F = 2.298, R2 = 0.103, p_adj_ = 0.003; weighted UniFrac Adonis, df = 1, F = 4.470, R2 = 0.183, p_adj_ = 0.015). Interestingly, at T1, no significant differences in beta diversity metrics of coral-associated bacterial communities could be found when comparing the two different priming methods employed (LP vs. DP: unweighted UniFrac Adonis, df = 1, F = 1.073, R2 = 0.060, p_adj_ = 0.254; weighted UniFrac Adonis, df = 1, F = 1.065, R2 = 0.059, p_adj_ = 0.468). Moreover, at T1, Shannon diversity index differed significantly among treatments (Kruskal-Wallis, χ² = 13.50, df = 3, p < 0.01; Supplementary Tables 6, 7), with pairwise comparisons indicating that both LP and DP samples exhibited significantly higher diversity than C and NP groups (Dunn’s test, p < 0.01), while no significant difference was observed between LP and DP. Unexpectedly, no significant differences in the microbiomes structure were encountered across treatments in alpha and beta diversity measures at T2 (Fig. 2A; Supplementary Tables 5 to 7). Finally, analyses of multivariate dispersion revealed no significant differences in beta dispersion across timepoints within any treatment, indicating that the observed treatment and time dependent differences reflected shifts in community composition rather than changes in within-group variability.

**Figure 2.**
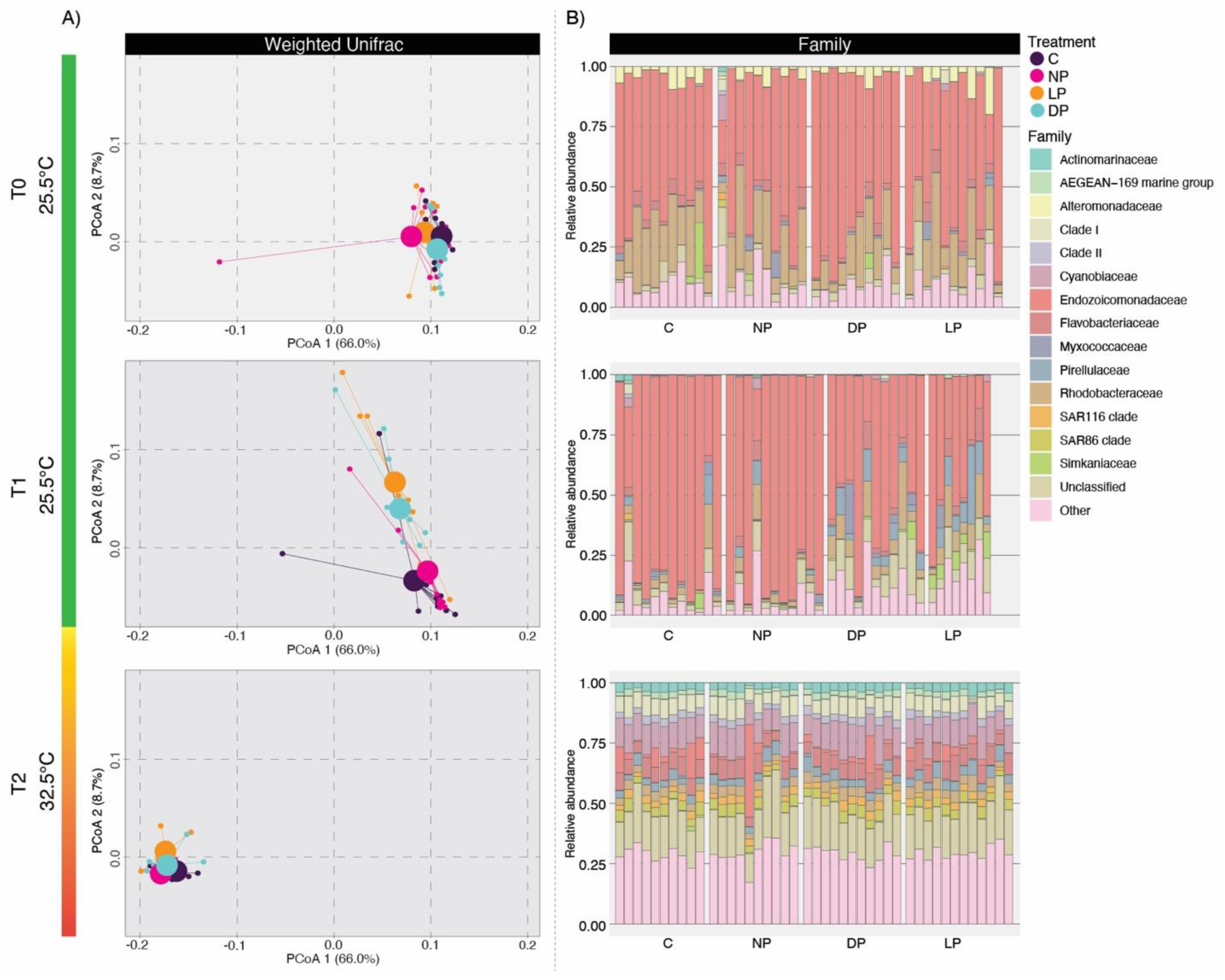
Dynamics of host-associated bacterial communities and composition shifts induced by priming and subsequent infection at increasing temperature. (A) PCoA graphs of weighted UniFrac distance matrix showing separation into clusters based on treatment group (C = control samples; NP = non-primed samples; LP = live VBAA450 primed samples; DP = inactivated VBAA450 primed samples). Small circles reflect communities within individual *P. favosa* fragments while large circles represent the centroids of the groups. (B) Barplots show mean relative abundance for the 15 most abundant bacterial families grouped by treatment at the different timeframes of the experiment. All remaining families were collapsed into “Other” category.

At T0, two families dominated the microbiome of *P. favosa*’s samples: Endozoicomonadaceae and Rhodobacteraceae, whose average relative abundances ranged between 52.002 % (± 4.142) – 67.335 % (± 5.290) and 14.515 % (± 2.881) – 20.646 % (± 3.103) of the reads regardless of treatments, respectively. The same two families were the most abundant also after priming, except in the LP treatments where the Pirellulaceae family followed Endozoicomonadaceae with a relative abundance of 10.526 % (± 3.472) (Fig. 2B, Supplementary Table 8). After heat stress and pathogen infection (T2), a dramatic change in bacterial community composition involved all treatments with the previously dominant Endozoicomonadaceae dropping to an average of 5.429 % (± 1.527) in control samples, 4.991 % ± (3.725), 2.505 % (± 0.560) and 2.869 % (± 1.282) in NP, LP and DP treatments respectively, while the families Cyanobiaceae and Flavobacteriaceae accounted for approximately the 11 – 12 % and 6 – 8 % respectively of the total prokaryotic reads in all treatments regardless of the priming strategy.

At the ASV level, the increase of ASVs in LP and DP groups identified as VBAA450 at 100 % identity over 420 bp (99% coverage of the V3 – V4 regions) with the deposited reference sequence confirmed its incorporation in the primed coral samples at T1. Particularly, Kruskal-Wallis test revealed a significant overall difference in VBAA450 read abundances among treatments (χ² = 30.2, df = 3, p < 0.001) with both priming treatments averaging a significantly higher relative abundance of VBAA450 compared with C and NP groups (T1: C_VBAA450_ = 0.001 % ± 0.0009; NP_VBAA450_ = 0.0005 % ± 0.0002; LP _VBAA450_ = 0.178 % ± 0.051; DP _VBAA450_ = 0.043 ± 0.038; Dunn’s test LP vs. C and NP: p*_adj_*. < 0.001; DP vs. C and NP: p*_adj_*. < 0.05). Surprisingly, at T2, despite the higher load of VBAA450 infection compared to the previous priming dosage (i.e., 10^4^ cells/ml final concentration for priming vs. 10^6^ cells/ml final concentration for subsequent infection), no significant differences were found among treatments after 15 days at increasing temperatures, although non primed samples hosted the highest relative abundance of pathogen reads compared to control and primed groups (T2: C_VBAA450_ = 0.002 % ± 0.001; NP_VBAA450_ = 0.007 % ± 0.003; LP _VBAA450_ = 0.002 % ± 0.0007; DP _VBAA450_ = 0.001 % ± 0.001).

Finally, ANCOM-BC2 differential abundance analysis indicated the presence of a high number of enriched bacterial ASVs between samples that received the two priming treatments compared to the C and NP groups after the priming phase at T1. When comparing LP samples to C and NP groups, a total of n = 564 (n = 179 enriched and n = 385 decreased) and n = 367 (n = 166 enriched and n = 201 decreased) ASVs were differentially abundant, respectively. A similar scenario was observed in samples primed with chemically inactivated VBAA450 (DP group) in which a total of n = 307 (n = 146 enriched and n = 161 decreased) and n = 354 (n = 205 enriched and n = 149 decreased) resulted significantly differentially abundant compared to C and NP controls respectively. Among the top 50 significantly differentially abundant ASVs in the LP and DP groups compared to the C and NP groups, 7 enriched and 20 decreased were common to the primed corals regardless of the priming strategy. Interestingly, significantly enriched ASVs in both LP and DP samples included validated and putative beneficial genera members of the families Rhodobacteraceae (i.e., *Ruegeria*), Halomonadaceae (i.e., *Halomonas* and *Cobetia*), Pseudoalteromonadaceae (i.e., *Pseudoalteromonas*), and Myxococcaceae (i.e., *P3OB-42*) (Fig. 3A, B and Supplementary Table 9). Conversely, only one ASV (i.e., Unclassified ASV 157 belonging to the phylum Patescibacteria) was differentially abundant when comparing non primed groups, as expected for samples that did not receive any treatment yet, while no ASV was detected as significantly enriched or decreased in the comparison between samples primed with live and chemically inactivated VBAA450 pathogen. At T2, reinforcing the non-significant results observed in alpha and beta diversity results, very few ASVs were detected as differentially abundant ranging from a maximum of 14 and a minimum of 1 ASVs when comparing NP with LP and NP with DP groups respectively. A complete list of the differentially abundant ASVs resulting from ANCOM-BC2 analyses (including unclassified ASVs) are detailed in Supplementary Table 9.

**Figure 3.**
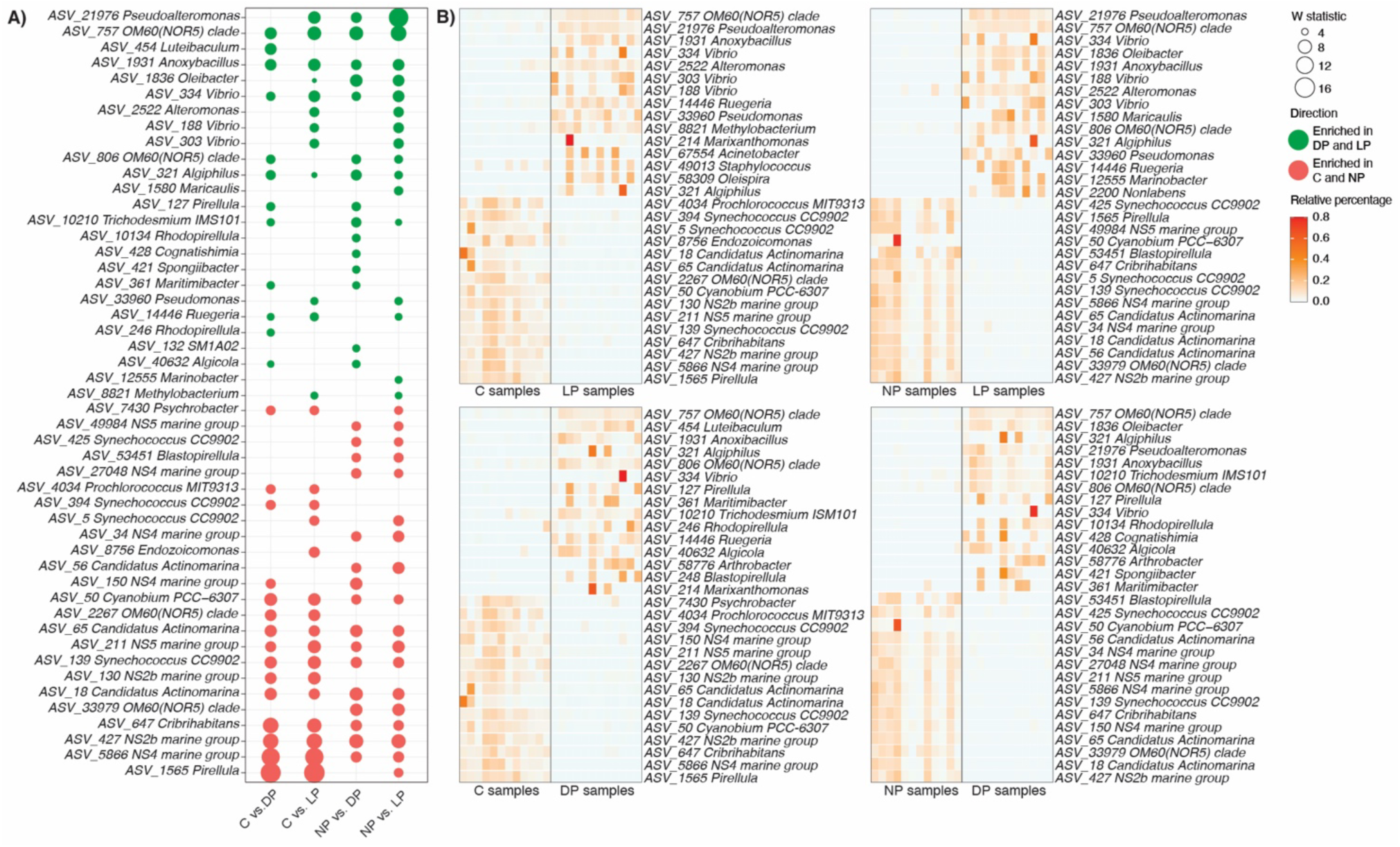
Differentially abundant ASVs after the priming regimen at T1. (A) Dotplot representing the overall top 50 differentially abundant ASVs with their genus taxonomic identification across comparisons between primed (LP and DP) and non-primed (C and NP) treatment groups. The complete lists of the differentially abundant ASVs resulting from ANCOM-BC2 analyses (including “Unclassified” ASVs) are available in Supplementary Table 9. (B) Heatmaps represent the relative percentage of the top 15 most enriched (top part) and 15 most decreased (bottom part) ASVs specifically for each pairwise comparison between primed vs. non-primed corals. The X-axis represents individual coral replicates for each treatment, respectively.

### Priming modulates distinct host gene expression pathways and responses

To investigate how prior microbial exposure influenced the host’s transcriptional response to subsequent thermal and pathogenic stress, we conducted host RNA sequencing across all four experimental treatments over the course of the experiment (4 conditions x 3 timepoints x 4 coral replicates = 48 samples). At T0, all samples clustered together, with minimal separation between the centroids of the treatment groups, indicating a consistent transcriptional state prior to the application of priming with no significant differences among any of the pairwise contrasts between treatments. By T1, transcriptomic profiles began to diverge particularly for the LP and DP treatments compared to the non-primed groups, suggesting a transcriptional response following priming with VBAA450, although not accompanied by statistical significance. At T2, following exposure to both heat and high VBAA450 infection load, samples from all challenged groups (i.e. NP, LP, DP) diverged from the controls (C) with NP corals showing the most substantial displacement compared to control samples even if not statistically significant (Fig. 4A). Nevertheless, when comparing treatments across the entire timeframe of the experiment principal component analysis (PCA) of transcriptomic profiles revealed overall significant changes in gene expression among the four experimental treatments (Adonis _(treatment+sampling_time)_, R2 = 0.047, df = 3, F = 0.849, Pr (>F) = 0.012). Specifically, significant differences in all pairwise contrasts were found, confirming an adaptive response by the host to priming and subsequent combined stressors: C vs. NP samples (Adonis _(timepoints)_, df = 2, R2 = 0.191, F = 2.501, Pr (>F) = 0.0001), C vs. DP samples (Adonis _(timepoints)_, df = 2, R2 = 0.186, F = 2.353, Pr (>F) = 0.001), C vs. LP samples (Adonis _(timepoints)_, df = 2, R2 = 0.198, F = 2.547, Pr (>F) = 0.001), NP vs. DP samples (Adonis _(timepoints)_, df = 2, R2 = 0.219, F = 2.980, Pr (>F) < 0.0001), NP vs. LP samples (Adonis _(timepoints)_, df = 2, R2 = 0.223, F = 3.016, Pr (>F) < 0.0001 ), and finally the two primed treatment groups (Adonis _(timepoints)_, df = 2, R2 = 0.229, F = 3.053, Pr (>F) = 0.0002) (Fig. 4A; Supplementary Table 10).

**Figure 4.**
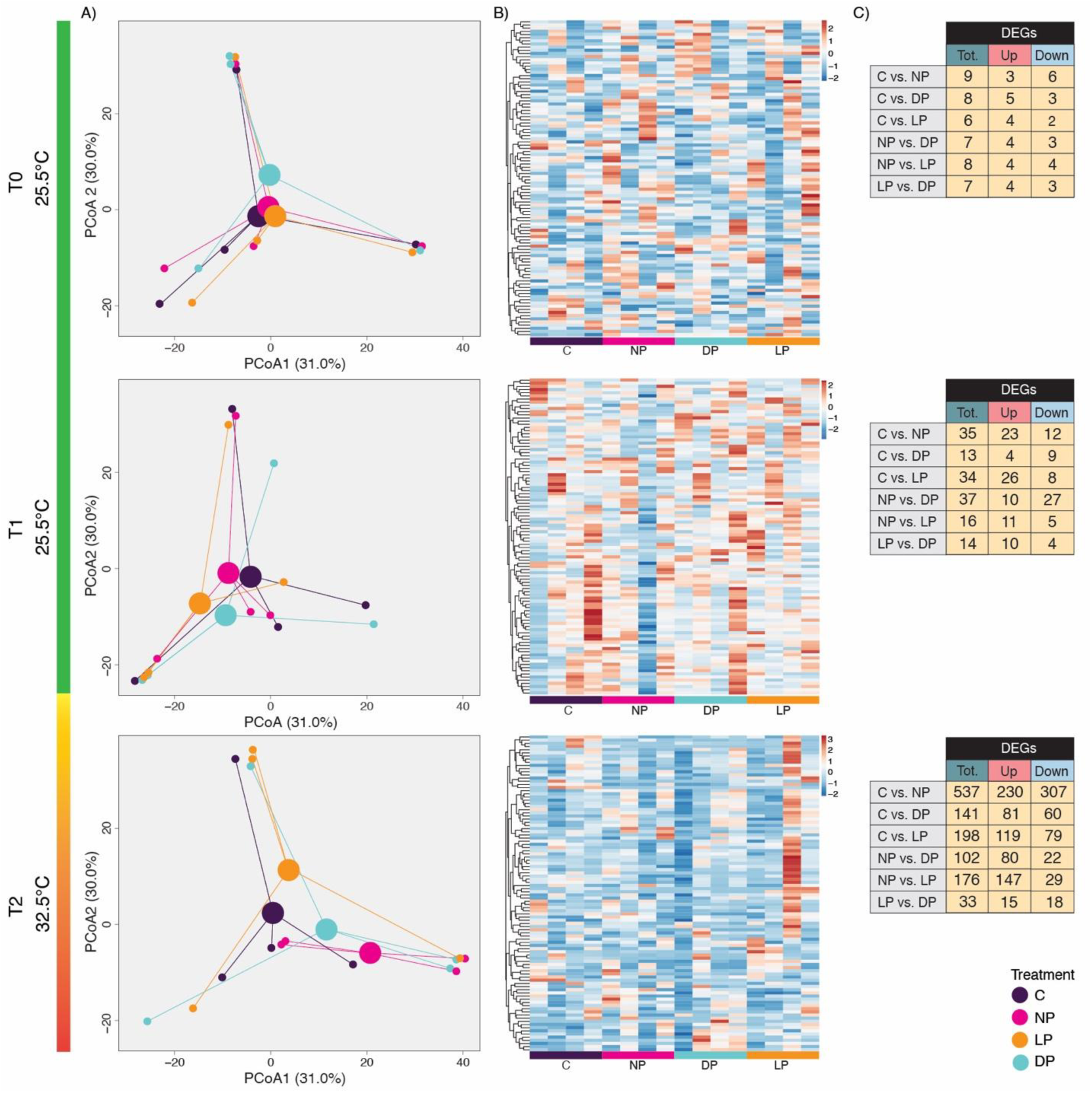
Coral host transcriptome shifts induced by priming and subsequent infection at increasing temperature. (A) PCoA graphs showing clustering and separation between clusters based on treatment groups (C = control samples; NP = non-primed samples; LP = live VBAA450 primed samples; DP = inactivated VBAA450 primed samples). Small circles reflect transcriptomic profiles associated with individual *P. favosa fragments* whereas large circles represent the centroids of the groups. (B) Heatmaps of the top 100 most variable expressed genes at the different time points in each sample. (C) Tables list the number of DEGS between pairwise treatment contrasts at the different timepoints of the study.

In line with the pairwise differences, we found distinct numbers of differentially expressed genes (DEGs) between treatments for the different timepoints (Fig. 4C; Supplementary Table 11).

Specifically, at T0, the number of DEGs across treatment comparisons was minimal, with no contrast exceeding nine DEGs. At T1, the number of DEGs observed in both primed groups compared with controls increased to a minimum of 13 (4 up and 9 down) between C and DP groups and a maximum of 37 DEGs (10 up and 27 down) between NP and DP groups. At T2, the number of DEGs dramatically increased with the contrast among controls and non-primed but infected samples, yielding the highest number (537 DEGs, 230 up and 307 down), followed by C vs. LP (198 DEGs, 119 up and 79 down) and C vs. DP (141 DEGs, 81 up and 60 down).

Conversely, comparison between primed groups and NP treatment revealed fewer DEGs, with a total of 176 (147 up and 29 down) and 102 (80 up and 22 down) between NP vs. LP and NP vs. DP treatments, respectively. Complete lists of differentially expressed genes for all pairwise treatment comparisons at each timepoint, along with the 15 most strongly up- and down-regulated transcripts per timepoint, are provided in Supplementary Table 11 and summarized visually in Supplementary Fig. 2 respectively.

Gene ontology (GO) enrichment analysis of the *P. verrucosa* host DEGs revealed distinct biological processes (BPs) in response to priming, heat stress, and high load pathogen infection across the timeframe of the experiment among treatments, reflecting biological processes associated with the baseline-to-outcome transcriptional shift over the course of the study (Fig. 5; Supplementary Table 12). Significantly enriched GO terms were summarized into 25 non-redundant GO clusters in C, 15 in NP, 14 in DP and 34 in LP samples by REVIGO analysis.

**Figure 5.**
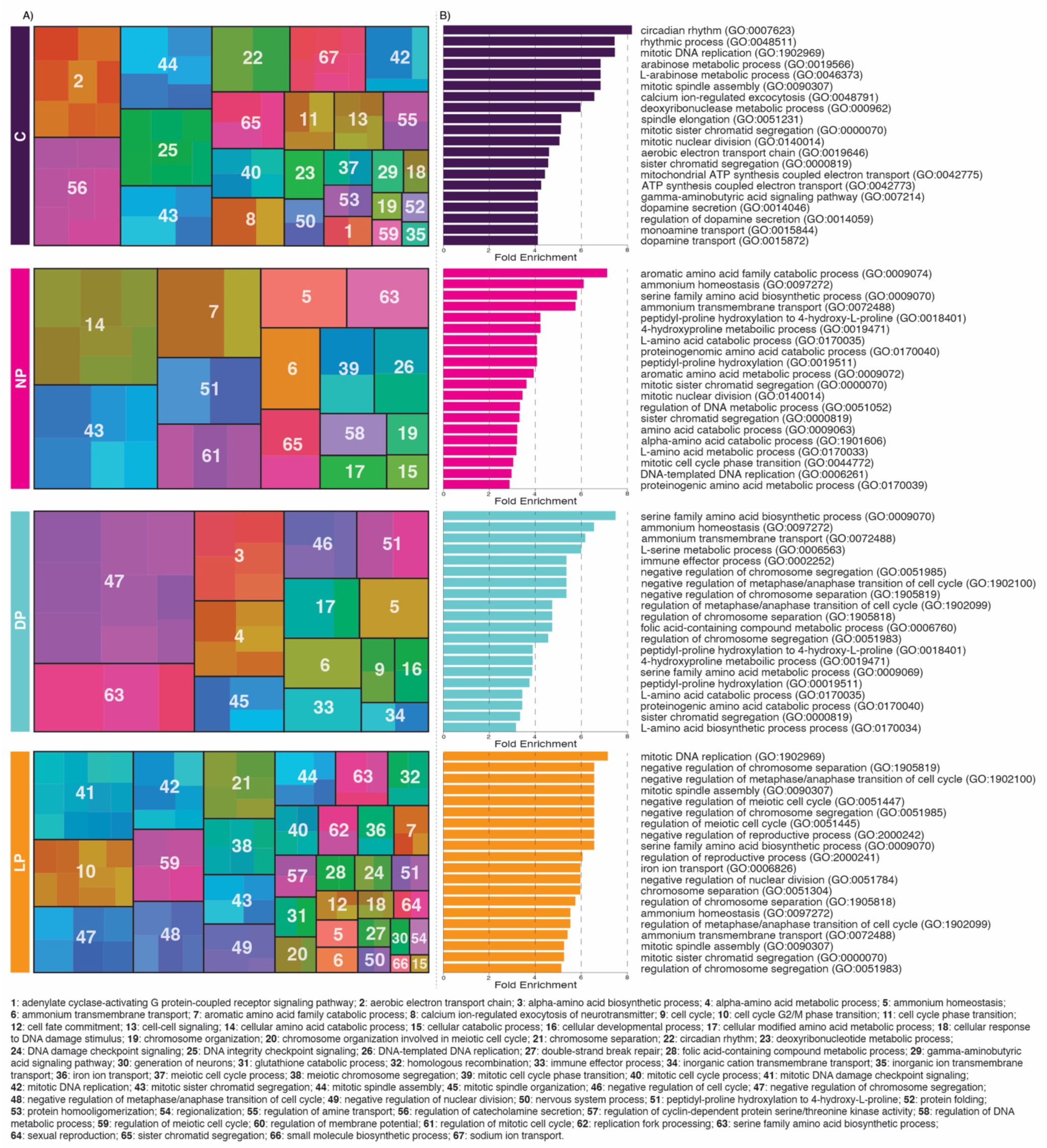
Gene ontology (GO) enrichment analysis of the *P. verrucosa* host DEGs through the timeframe of the experiment. (A) Comparison of significantly enriched GO terms visualized as treemaps where semantically redundant GO terms were reduced using REVIGO for each treatment (C = control samples; NP = non-primed samples; LP = live VBAA450 primed samples; DP = inactivated VBAA450 primed samples). Each colored rectangle represents a non-redundant biological process GO term. The size of each rectangle reflects the Fold Enrichment of the corresponding GO term, while color groups similar biological functions into clusters delimited by black boxes. White labels display representative GO terms selected by REVIGO to summarize each cluster, prioritizing terms with high relevance and lower redundancy. (B) Bar plots panel listing the top 20 GO terms significantly enriched (p_adj._ < 0.05) per treatment ranked by Fold Enrichment. The full list of GO terms is listed in Supplementary Table 12.

More in detail the C group, as response to only heat stress, showed significant enrichment (p*_adj._* < 0.05) of BPs associated with a broad stress response and redox homeostasis, metabolism and energy rewiring, and with nuclear and chromosome dynamics, including ‘L-arabinose metabolic process’ (GO:0046373), ‘cellular response to stress’ (GO:0033554), and ‘mitotic nuclear division’ (GO:0140014). Conversely, non-primed corals exposed to pathogen infection (NP) exhibited a transcriptional profile dominated by terms suggesting metabolic imbalance and stress adaptation failure under combined heat and pathogen stressors, such as upregulation of genes linked to ‘aromatic amino acid family catabolic process’ (GO:0009074), ‘L-amino acid catabolic process’ (GO:0170035) and downregulation of genes associated with ‘ammonium homeostasis’ (GO:0097272) and ‘peptidyl-proline hydroxylation’ (GO:0019511), which reflect enhanced amino acid breakdown for emergency and redox balance together with impaired regulation of nitrogen homeostasis and protein modification.

Regarding the two primed treatments, samples from DP and LP groups showed differences in the enriched host encoded BPs. For example, in DP corals enriched BPs included amino acid biosynthesis and nitrogen regulation, alongside immune effector activity [‘immune effector process’ (GO:0002252)]. Notably, several complement immune pathway components, including but not limited to ‘complement factor B-like’ (LOC131771060; log_2_FoldChange = 3.055, p*_adj_*. = 0.022), and ‘complement C2’ (LOC131775549; log_2_FoldChange = 3.985, p*_adj_*. < 0.001), were strongly upregulated. Conversely, LP samples showed a pronounced emphasis on mitotic and genomic maintenance pathways evidenced by BPs largely involving downregulated genes.

Despite the absence of complement-related GO significant enrichment, several genes linked to the complement immune pathway were upregulated in live-primed samples [e.g., C2-like’ (LOC136280257; log_2_FoldChange = 1.811, p*_adj_*. = 0.013) and ‘complement C2’ (LOC131775549; log_2_FoldChange = 1.529, p*_adj_*. = 0.010)], whereas complement C1q-like protein 4’ was significantly downregulated (LOC131782126; log_2_FoldChange = - 4.885, p*_adj_*. < 0.001), paralleling in part the DP response. Both LP and DP corals exhibited broad transcriptional activation of TNF receptor-associated factors (TRAFs) genes, with LP group showing marked immune activation compared to NP samples. For instance, ‘TNF receptor-associated factor 6 (TRAF-6; LOC131787738) was massively upregulated (log_2_FoldChange = 6.474, p*_adj_*. = 0.023), representing one of the strongest transcriptional responses observed. Several leucine-rich repeat (LRR) domain-containing genes, encoding for pattern-recognition receptors (PRRs) mediating the recognition of pathogen-associated patterns (PAMPs), were also upregulated, such as LOC131773716 (log_2_FoldChange = 1.064, p*_adj_*. = 0.003), LOC131799288 (log_2_FoldChange = 3.635, p*_adj_*. = 0.049) and LOC131790242 (log_2_FoldChange = 0.997, p*_adj_*. = 0.011). Notably two genes associated with inflammasome-like signaling were significantly downregulated in LP corals compared to the non-primed group: the CARD- and ANK-domain containing inflammasome adapter protein-like (LOC131772013, log_2_FoldChange = -3.639, p*_adj_*. = 0.007) and NLR family CARD domain-containing protein 3-like (LOC131785002, log_2_FoldChange = - 2.982, p*_adj_*. = 0.030). This pattern may reflect a more refined immune response following priming with live VBAA450. The complete list of DEGs resulting from DESeq2 differential gene expression analysis and GO terms are detailed in Supplementary Tables 11 and 12.

## DISCUSSION

Here we provide a proof of principle that the coral holobiont can be primed by both live and chemically inactivated pathogens to increase resilience in subsequent exposures by shifting the assemblage of the coral-associated bacterial communities and altering the coral host’s transcriptomic response. Studies on other marine invertebrates revealed similar patterns. For example, a study conducted by Yang et al. (92) showed the efficacy of immune priming against *Vibrio parahaemolyticus* in individuals of the mud crab species *Scylla paramamosain*.

Particularly, specimens that received a priming dosage of 2 x 10^5^ cells/g wet weight of formalin-inactivated pathogen demonstrated approximately a 75 % higher survival rate than unprimed individuals after being challenged with live *V. parahaemolyticus*. Similarly, in Cnidarians, Brown and Rodriguez-Lanetty (43) provided evidence that *Exaiptasia pallida* anemones, previously exposed to *V. coralliilyticus* under sub-lethal conditions, exhibited a significantly increased survivorship compared to naïve samples when subjected to a subsequent lethal challenge. Here, we specifically show that priming *P. favosa* with *V. coralliilyticus* induces significant shifts in the host-associated bacterial communities without compromising coral health, as evidenced by the lack of significant differences in pathogen induced bleaching and photochemical efficiency of the coral associated Symbiodiniaceae. This finding suggests that, the observed microbiome shifts may reflect an adaptive response indicative of reactive homeostasis and/or a healthy defensive state rather than indicating dysbiosis as a response to sublethal doses of either live or chemically inactivated *V. coralliilyticus* (93,94). Interestingly primed coral microbiomes converged towards a similar configuration regardless of the priming strategy adopted. This likely reflects the use of chemical inactivation, rather than heat-killing methods, preserving critical features for host immune recognition and microbiome modulation. To the best of our knowledge, no direct study is present in the scientific literature comparing the responses to chemically inactivated versus heat-killed *Vibrio coralliilyticus* in coral models. However, consistent patterns have emerged from studies on other *Vibrio* species in aquatic organisms that highlight the limitations of heat-killing and the advantages of chemical inactivation. For example, heat-killing methods are known to denature key microbial-associated molecular patterns (MAMPs), degrade outer membrane vesicles (OMVs) and surface proteins leading to muted reactive responses due to the loss of key immunostimulatory features. In contrast, chemical inactivation, such as the use of formalin in our case, has been shown to preserve integrity of surface-exposed antigens resulting in a stronger immune recognition, microbiome modulation and better mimicry of responses to live pathogen infection (95–97) Moreover, both live and chemically inactivation priming strategies resulted in an enrichment of the coral microbiome in bacterial ASVs belonging to putative, as well as previously validated, beneficial genera including *Ruegeria*, *Halomonas*, *Cobetia* and *Pseudoalteromonas* (65,98–100), although functional roles can vary at the species level and such genus-level assignments should be interpreted with caution. Among these, members of the genus *Ruegeria* have been shown to inhibit the growth of *V. coralliilyticus* and to mitigate bleaching symptoms induced by pathogen infection (101,102). Similarly, several *Pseudoalteromonas* strains are recognized as coral probiotics because of their ability to produce antimicrobial compounds and enhance coral resistance to both heat stress and disease (65,98–100). This selective enrichment in beneficial bacterial taxa aligns with the coral probiotic hypothesis and adaptive microbiome hypothesis, which suggest that exposure to stressors (such as a sub-lethal pathogen doses) can prime the host-associated microbial communities fostering beneficial members which are able to quickly proliferate and/or activate the production of defensive compounds (11,103). This phenomenon has been observed not only in corals but also in other organisms, such as amphibians, where microbial restructuring following mild stress or infection reduced susceptibility to future pathogen exposures thanks to the enrichment in beneficial prokaryotes (94,104,105).

Nevertheless, in our study, after 15 days of exposure to high temperatures (T2), the bacterial communities of all samples converged regardless of the treatments applied. This pattern indicates that prolonged thermal stress acted as a major driver of microbiome structure at the terminal time point, reducing between-treatments differences and leading to increased similarity across communities. Under these conditions, any effects of high load pathogen infection occurred within a microbiome strongly constrained by temperature and therefore did not manifest as distinct treatment-specific taxonomic patterns. Such convergence is consistent with the well-established role of temperature as a dominant driver of marine host-associated microbiome composition, often linked to the emergence of generalist microbial taxa and the decline of specialized symbionts (106–108). In this context, the priming-associated microbiome shifts observed at T1, such as the enrichment of beneficial taxa, were no longer maintained as distinct taxonomic signatures under prolonged thermal and pathogenic stress. In our study, the observed increase in alpha diversity metrics at T2 across all treatments, may indicate a shift toward a broader functional microbiome portfolio, consistent with similar findings in other coral species and marine sponges (109,110). Despite this taxonomic homogenization across treatments observed at T2, the significantly higher photosynthetic efficiency coupled with a lower *Vibrio*-induced bleaching recorded in the primed groups may be evidence that past exposure to the pathogen conditioned microbial responses and had a lasting functional effect on the holobiont system. This suggests that, although the bacteria communities appeared similar in taxonomic composition at T2, they may have retained functionally distinct capabilities in the primed corals. In other words, microbiome shifts following priming led to functional changes that were not reflected in community structure alone. This aligns with the growing recognition that microbiomes can undergo functional reprogramming and modulate their roles in response to previous stress, allowing their host to maintain physiological advantages even when community composition appears convergent (111,112).

In accordance with our results, recent research supports the hypothesis of ecological memory within the coral holobiont, showing that past disturbances (e.g., heat weaves) can induce microbial acclimatization and induce increased resistance to repeated stress events (10,113). It is then tempting to speculate that a similar form of microbiome-mediated memory may protect coral holobionts against recurrent pathogen exposure by maintaining or reactivating functional microbial traits that support host resilience. However, caution is warranted when interpreting the results of our study as it is important to differentiate between ‘short-term microbial conditioning’ and long-term microbiome memory especially considering the short four-day rest period between priming and subsequent challenge. Microbiome memory, as traditionally defined, tends to unfold over extended timescales (weeks or months) or even across generational scales (10,114). In contrast, the rapid timeline of our experiment between the priming phase and subsequent infection suggests that the enhanced resilience observed in primed corals could be more accurately described as ‘microbial conditioning’ or ‘microbiome training’, wherein rapid transcriptional reprogramming and metabolic priming enable faster and more robust response to repeated stress by potentially optimizing antibacterial compounds production, antioxidant production, and nutrient cycling. Such rapid priming mechanisms have been previously observed in other marine organisms, for example in oysters and seagrasses, where exposure to low-dose stressors has been shown to trigger immediate microbial reconfiguration, enhancing survival during subsequent environmental challenges (115,116).

While such microbiome training likely contributes to coral resilience, it may act in concert with host-level regulatory processes. In recent years, the concept of immunological priming has expanded beyond vertebrates, challenging the long-standing paradigm that invertebrates, such as corals, rely solely on innate responses. Increasing evidence supports the existence of antigen-independent immunological memory in invertebrates, called ‘trained immunity’, which show that prior exposure to microbial stimuli can reprogram the host immune system via transcriptional, metabolic and possibly epigenetic mechanisms (117,118). Accordingly, our coral transcriptome analysis revealed a dynamic reconfiguration of host gene expression across treatments and time, reflecting complex interactions between priming history and subsequent exposure to paired thermal and pathogenic stressors. Because pathogen priming is inherently a history-dependent process, whose effects are defined not solely by the terminal transcriptional state but by how prior exposure reshapes the trajectory of the host response to subsequent stress, we emphasized within-treatment longitudinal comparisons across the full experimental timeline rather than cross-treatment contrasts at individual time points, thereby enabling a more robust assessment of priming-induced transcriptional reprogramming and its consequences for biological process enrichment at the whole-experiment scale.

After coral acclimation (T0), coral transcriptomes clustered across all treatments, indicating a stable baseline between groups prior to experimental manipulation. By T1, transcriptomic divergence began to emerge for the primed corals compared to controls and non-primed samples, while following thermal stress and high dose pathogen challenge (T2), all exposed treatments diverged from the uninfected control with corals primed with either live or inactivated *V. coralliilyticus* displaying transcriptomic profiles indicative of a trained immunity-like state. Despite facing the same high pathogen load and high temperature regimes, both primed groups exhibited fewer differentially expressed genes than non-primed corals compared to controls. This may indicate a form of transcriptional modulation, often described as transcriptional buffering, where prior exposure tempers widespread transcriptional upheaval during subsequent stress events, enabling the host to activate a more refined and targeted gene expression instead of a broader but more dysregulated one (119,120). Moreover, beyond DEGs count, NP corals showed upregulation in biological processes associated with physiological breakdown. Specifically, non-primed samples were enriched for catabolic and stress-associated biological processes, such as ‘aromatic amino acid catabolic process’ and ‘oxoacid metabolic process’ coupled with downregulation of homeostatic functions indicative of metabolic exhaustion. Conversely, the gene expression profile of LP corals was distinctive, with gene ontology enrichment analysis revealing a strong bias toward regulation of cell cycle and mitotic checkpoints characterized by downregulated GO terms such as ‘mitotic DNA replication’, ‘chromosome segregation’, ‘organelle fission’ and ‘nuclear division’. These transcriptomic restraints are consistent with previous findings where corals exposed to chronic stressors exhibit improved performance under subsequent acute stress, potentially due to conserved acclimatory or adaptive cellular mechanisms (121,122). Repression of cell cycle related processes and maintenance of genomic integrity are thought to safeguard cellular homeostasis allowing corals to prioritize repair and defense over proliferation during stress which mimic observations in vertebrate systems where trained immunity is often accompanied by metabolic rewiring and chromatin remodeling that dampen unnecessary responses while amplifying specific effector functions (117).

At the immune signaling level, both primed groups exhibited broad transcriptional activation of tumor necrosis factor (TNF) receptor-associated factors (TRAFs). Among these, TRAF6-like was significantly upregulated in LP versus NP corals. In vertebrates, TRAF6 is a central adaptor protein in toll-like receptor (TLR) and TNF receptor signaling pathways that mediates the activation of downstream cascades such as NF-kB and MAPKs, promoting immune readiness and cell survival (123). Different studies in cnidarians strongly support the presence and evolutionary conservation of these pathways (124,125) with NF-kB signaling that has been shown to be functionally responsive to immune challenge in *Orbicella faveolata*, reinforcing its relevance in coral immunity (126). Other TRAFs, also differentially expressed in the primed groups, likely contribute to a fine-tuned regulation of both immune activation and feedback control, enabling primed corals to modulate inflammatory signaling with greater precision rather than necessarily increasing inflammatory tone (127,128).

In parallel, both DP and LP corals exhibited differential regulation of complement system components, indicating an additional layer of immune modulation linked to priming. This differential expression suggests a strategic tuning of upstream recognition components, potentially enabling the host to optimize microbial clearance while minimizing collateral immune-mediated damage. LP corals also showed a significant downregulation of key inflammasome-associated genes when compared to non-primed samples, including a CARD- and ANK-domain containing inflammasome adapter protein-like and NLR family CARD domain-containing protein, which have been proposed to mediate pro-inflammatory cytokine release and pyroptosis as seen in vertebrates (129,130). Their significant downregulation in live-primed corals compared to non-primed ones may be indicative of a dampened inflammatory potential, possibly as a protective mechanism to avoid immune-mediate tissue damage.

The coordinated upregulation of TRAF signaling pathways, selective engagement of complement factors, and suppression of inflammasome effectors in primed corals, particularly in the LP group, support the emergence of a strategically modulated trained immunity-like phenotype, key for immunological balance, which may maintain readiness for microbial recognition while limiting immunological overactivation. Altogether our host transcriptomic results point to the fact that primed corals appear to engage in a layered immune and modulated cellular mechanism strategy, optimizing both surveillance and self-preservation under stress compared to the seemingly uncoordinated response observed in non-primed samples.

While our study provides insights into the interplay between microbial priming and coral holobiont response, we need to acknowledge that the relatively short interval between priming and subsequent pathogen challenge constrains our ability to evaluate the long-term persistence of the observed priming effects. Future studies combining extended resting periods and finer temporal sampling would be crucial to disentangle holobiont transient conditioning from a potential more stable microbiome and host immune memory. Moreover, although our gene expression data suggest the presence of trained immunity-like features, yet we did not assess metabolic or epigenetic changes that often underlie this phenomenon. Finally, ‘genotype’ likely represents a source of biological variation that was not explicitly modeled in the present analysis. Although our balanced experimental design ensured that coral fragments from the same colonies were distributed across treatments, timepoints and tank replicates, future studies with increased colony replication would enable population-level inference and allow ‘genotype’ to be modeled explicitly, for example as a random effect within mixed-model frameworks (e.g., Dream (131)).

Nevertheless, taken altogether, our findings challenge the long-held assumption that coral rely only on innate mechanisms to cope with pathogen infections, contributing to the growing body of evidence questioning coral immunity as a static response (132). By integrating microbiome profiling and host transcriptomics, we show that microbial priming, whether with live or chemically inactivated *Vibrio coralliilyticus*, can influence both microbial community composition and host gene regulation supporting enhanced resistance to subsequent infection. These results highlight the coral holobiont as an interconnected and adaptable unit, capable of microbial and transcriptional reprogramming in response to priming and subsequent exposure to the same pathogen. Rather than relying solely on a reactive innate system, the *P. favosa* corals appear to leverage microbial cues to condition their responses potentially through a finely tuned crosstalk between host immunity and its associated bacterial communities enhancing resilience under compounded stress.

## DATA AVAILABILITY

Data and code are available upon publication and request.

## AUTHOR CONTRIBUTIONS

Conceptualization, M.M., N.G.B., F.C.G. and R.S.P.; methodology, M.M., N.G.B., F.C.G., E.P.S., A.G., M.C., G.A., K.S., C.P.A., L.C., L.B. and C.R.V.; investigation, M.M., N.G.B., F.C.G., E.P.S., A.G., M.C., G.A., K.S., C.P.A., L.C., L.B., C.R.V. and R.S.P., writing: original draft, M.M.; writing: review & editing, N.G.B., F.C.G., A.G., C.P.A., C.R.V. and R.S.P.; resources R.S.P. and C.R.V.

## ACNKOWLEDGEMENTS

The authors thank Zenon Batang, Nabeel M. Alikunhi, and the CMR team for allocation of workspace and technical support at the Core Lab for Coastal and Marine Resources (CMR) at KAUST.

