## Supplementary material for "Pathogen priming reveals host immune training and microbiome conditioning in corals"

**Supplementary materials**

Supplementary Table 8, 9, 11 and 12 (as .xlsx files) will be available upon publication at 10.5281/zenodo.17279049.

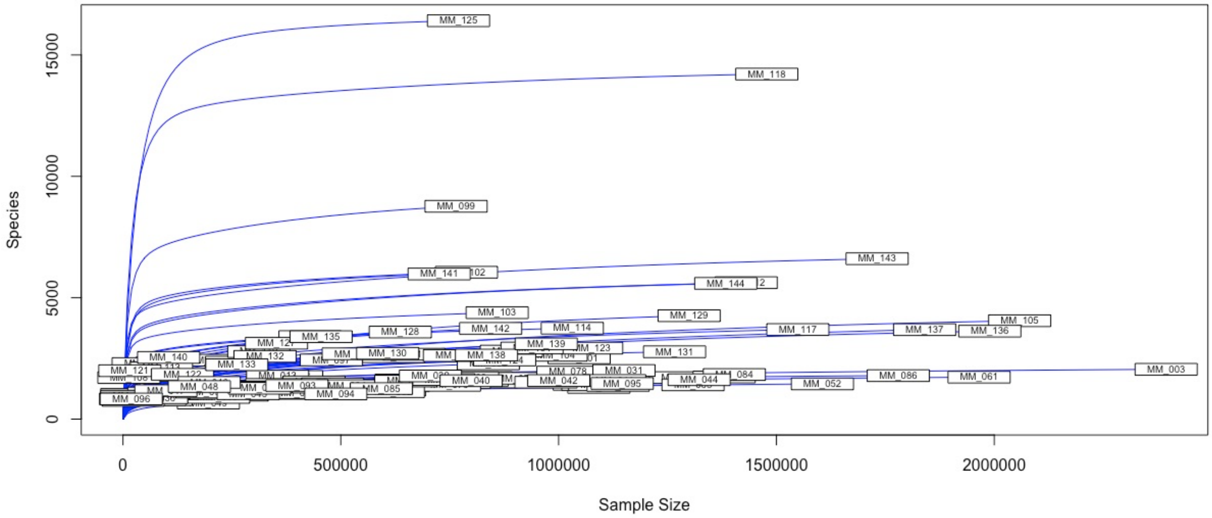


**Supplementary Figure 1.** Reads accumulation curve for microbiome analysis. Red-dashed line represents the 100,00 reads cutoff at which rarefaction was applied. X-axis represents sequencing depth while Y-axis represents the ASVs number.

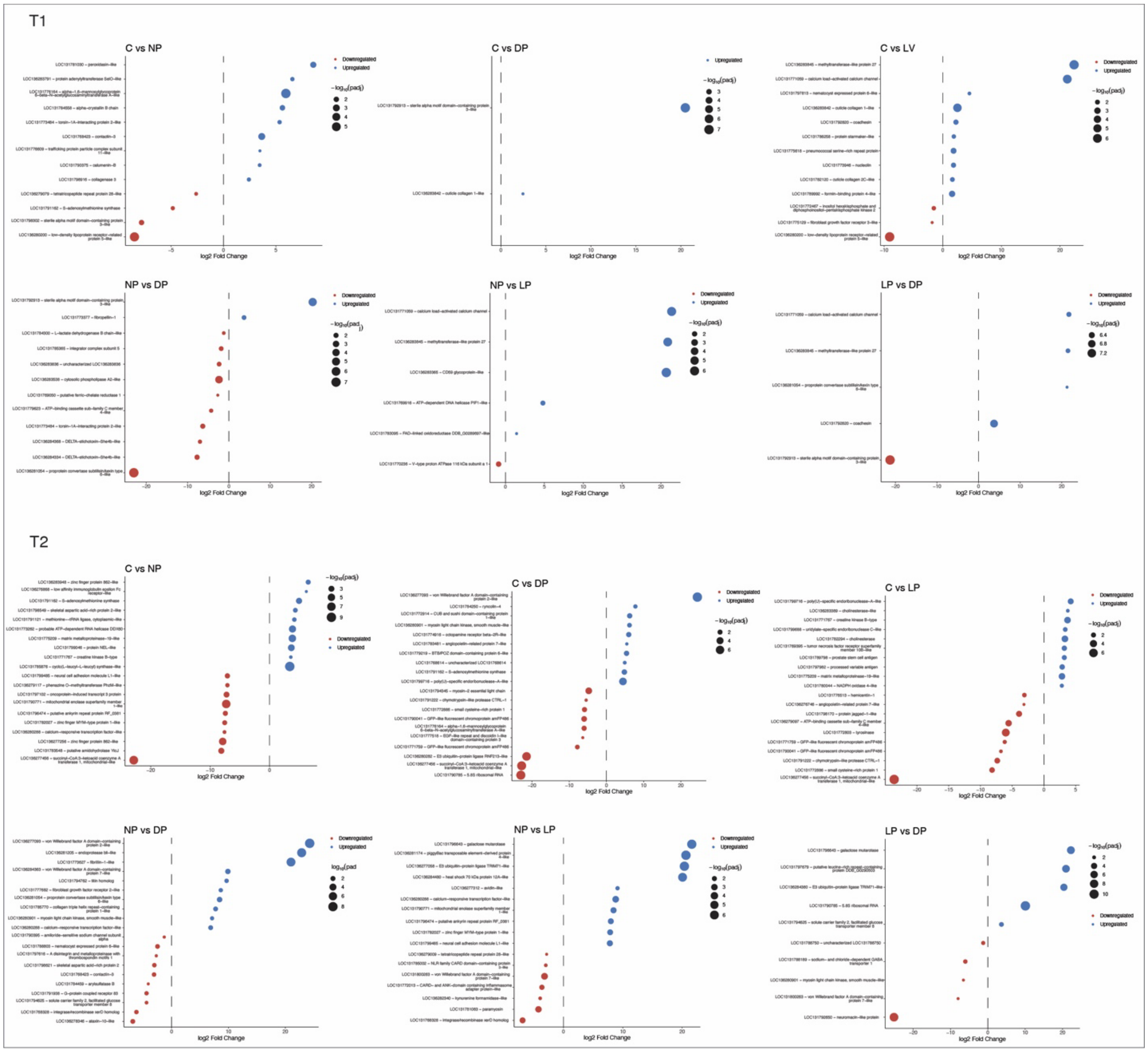


**Supplementary Figure 2**. Dotplots representing the 15 most significantly upregulated and 15 most significantly downregulated genes for each treatment pairwise comparison at T1 and T2.

**Supplementary Table 1.** Mean (± standard error) gray value representing the unweighted average brightness across all pixels within the coral fragment (from 0 = black to 255 = white) for each treatment at the reference temperatures.

| Treatment | Temperature (°C) | Mean gray value |
|---|---|---|
| C | 25.5 (T1) | 116.20 (±0.48) |
| NP | | 115.60 (±0.77) |
| DP | | 114.82 (±0.75) |
| LP | | 116.34 (±0.75) |
| C | 28.3 | 121.71 (±1.39) |
| NP | | 123.70 (±1.18) |
| DP | | 125.07 (±1.67) |
| LP | | 124.16 (±0.79) |
| C | 32.5 (T2) | 125.28 (±1.67) |
| NP | | 160.67 (±7.18) |
| DP | | 137.98 (±3.00) |
| LP | | 128.51 (±2.12) |

**Supplementary Table 2.** General linear mixed-effects models (GLMMs) with a 'Gamma' family distribution and with a 'log link' function on coral fragments' pixels unweighted average gray channel value**.**

Model_BL_T1_glmm

| | npar | AIC | BIC | logLik | deviance | Chisq | Df | Pr(>Chisq) |
|---|---|---|---|---|---|---|---|---|
| #Model_BL_T1_null | 4 | 503.84 | 514.10 | -247.92 | 495.84 | | | |
| #Model_BL_T1_glmm | 25 | 530.56 | 594.67 | -240.28 | 495.84 | 15.285 | 21 | 0.8084 |
| **Pairwise contrast** | | | | | | | | |
| contrast | estimate | SE | z.ratio | p.value | | | | |
| #C - NP | 1.005 | 0.00775 | 0.665 | 0.9101 | | | | |
| #C - DP | 1.012 | 0.00775 | 1.549 | 0.4081 | | | | |
| #C - LP | 0.999 | 0.00775 | -0.108 | 0.9996 | | | | |
| #NP - DP | 1.007 | 0.00775 | 0.884 | 0.8134 | | | | |
| #NP - LP | 0.994 | 0.00775 | -0.773 | 0.8666 | | | | |
| #DP - LP | 0.987 | 0.00775 | -1.657 | 0.3468 | | | | |

Model_BL_28_glmm

| | npar | AIC | BIC | logLik | deviance | Chisq | Df | Pr(>Chisq) |
|---|---|---|---|---|---|---|---|---|
| #Model_BL_28_null | 4 | 272.02 | 279.50 | -132.01 | 264.02 | | | |
| #Model_BL_28_glmm | 25 | 270.06 | 283.16 | -128.03 | 256.06 | 7.9571 | 3 | **0.04691** |
| **Pairwise contrast** | | | | | | | | |
| contrast | estimate | SE | z.ratio | p.value | | | | |
| #C - NP | 0.983 | 0.0109 | -1.521 | 0.4248 | | | | |
| #C - DP | 0.969 | 0.0108 | -2.813 | 0.2253 | | | | |
| #C - LP | 0.976 | 0.0109 | -2.157 | 0.1354 | | | | |
| #NP - DP | 0.986 | 0.0110 | -1.292 | 0.5680 | | | | |
| #NP - LP | 0.993 | 0.0111 | -0.636 | 0.9203 | | | | |
| #DP - LP | 1.007 | 0.0112 | 0.656 | 0.9135 | | | | |

Model_BL_T2_glmm

| | npar | AIC | BIC | logLik | deviance | Chisq | Df | Pr(>Chisq) |
|---|---|---|---|---|---|---|---|---|
| #Model_BL_T2_null | 4 | 420.25 | 427.74 | -206.12 | 412.25 | | | |
| #Model_BL_T2_glmm | 25 | 378.14 | 391.24 | -182.07 | 364.14 | 48.108 | 3 | **2.019e-10** |
| **Pairwise contrast** | | | | | | | | |
| contrast | estimate | SE | z.ratio | p.value | | | | |
| #C - NP | 0.783 | 0.0230 | -8.335 | **<.0001** | | | | |
| #C - DP | 0.909 | 0.0267 | -3.242 | **0.0065** | | | | |
| #C - LP | 0.975 | 0.0286 | -0.860 | 0.8253 | | | | |
| #NP - DP | 1.162 | 0.0341 | 5.096 | **<.0001** | | | | |
| #NP - LP | 1.246 | 0.0366 | 7.475 | **<.0001** | | | | |
| #DP - LP | 1.072 | 0.0315 | 2.382 | 0.0805 | | | | |

**Supplementary Table 3.** Average (± standard error) *Fv/Fm* values for each treatment at the reference temperatures.

| Treatment | Temperature (°C) | Fv/Fm |
|---|---|---|
| C | 25.5 (T0) | 0.621 (±0.007) |
| NP | | 0.624 (±0.002) |
| DP | | 0.621 (±0.002) |
| LP | | 0.631 (±0.003) |
| C | 25.5 (T1) | 0.627 (±0.004) |
| NP | | 0.629 (±0.002) |
| DP | | 0.622 (±0.002) |
| LP | | 0.628 (±0.005) |
| C | 28.3 | 0.622 (±0.006) |
| NP | | 0.599 (±0.006) |
| DP | | 0.622 (±0.005) |
| LP | | 0.622 (±0.005) |
| C | 31.1 | 0.588 (±0.006) |
| NP | | 0.538 (±0.012) |
| DP | | 0.581 (±0.008) |
| LP | | 0.584 (±0.005) |
| C | 32.5 (T2) | 0.589 (±0.006) |
| NP | | 0.443 (±0.032) |
| DP | | 0.565 (±0.005) |
| LP | | 0.577 (±0.007) |

**Supplementary Table 4.** General linear mixed-effects models (GLMMs) with a 'Gamma' family distribution and with a 'log link' function on coral *Fv/Fm* values.

Model_25T0_glmm

| | npar | AIC | BIC | logLik | deviance | Chisq | Df | Pr(>Chisq) |
|---|---|---|---|---|---|---|---|---|
| #Model_T0_null | 4 | -620.6 | -608.72 | 314.3 | -628.6 | | | |
| #Model_25T0_glmm | 7 | -617.71 | -596.93 | 315.86 | -631.71 | 3.1143 | 3 | 0.3743 |

**Pairwise contrast**

| contrast | estimate | SE | z.ratio | p.value |
|---|---|---|---|---|
| #C - NP | -0.00490 | 0.00998 | -0.491 | 0.9611 |
| #C - DP | -0.00013 | 0.00998 | -0.013 | 1 |
| #C - LP | -0.01538 | 0.00998 | -1.542 | 0.4126 |
| #NP - DP | 0.004778 | 0.00998 | 0.479 | 0.9638 |
| #NP - LP | -0.01048 | 0.00998 | -1.05 | 0.7198 |
| #DP - LP | -0.01526 | 0.00998 | -1.529 | 0.4201 |

Model_25T1_glmm

| | npar | AIC | BIC | logLik | deviance | Chisq | Df | Pr(>Chisq) |
|---|---|---|---|---|---|---|---|---|
| #Model_T1_null | 4 | -496.76 | -486.50 | 252.38 | -504.76 | | | |
| #Model_25T0_glmm | 7 | -493.14 | -475.19 | 253.57 | -507.14 | 2.3779 | 3 | 0.4978 |

**Pairwise contrast**

| contrast | estimate | SE | z.ratio | p.value |
|---|---|---|---|---|
| #C - NP | -0.00196 | 0.00781 | -0.251 | 0.9945 |
| #C - DP | 0.00909 | 0.00781 | 1.165 | 0.6493 |
| #C - LP | 0.00466 | 0.00781 | 0.597 | 0.9330 |
| #NP - DP | 0.01105 | 0.00781 | 1.415 | 0.4897 |
| #NP - LP | 0.00662 | 0.00781 | 0.847 | 0.8317 |
| #DP - LP | -0.00443 | 0.00781 | -0.568 | 0.9417 |

Model_28_glmm

| | npar | AIC | BIC | logLik | deviance | Chisq | Df | Pr(>Chisq) |
|---|---|---|---|---|---|---|---|---|
| #Model_28_null | 4 | -218.30 | -210.82 | 113.15 | -226.30 | | | |
| #Model_28_glmm | 7 | 0030 | -210.98 | 119.04 | -238.08 | 11.779 | 3 | **0.008181** |

**Pairwise contrast**

| contrast | estimate | SE | z.ratio | p.value |
|---|---|---|---|---|
| #C - NP | 0.038497 | 0.0127 | 3.041 | **0.0126** |
| #C - DP | 0.000498 | 0.0127 | 0.039 | 1.0000 |
| #C - LP | 0.000765 | 0.0127 | 0.060 | 0.9999 |
| #NP - DP | -0.03799 | 0.0127 | -3.001 | **0.0143** |
| #NP - LP | -0.03773 | 0.0127 | -2.980 | **0.0153** |
| #DP - LP | 0.00026 | 0.0127 | 0.021 | 1.0000 |

Model_31_glmm

| | npar | AIC | BIC | logLik | deviance | Chisq | Df | Pr(>Chisq) |
|---|---|---|---|---|---|---|---|---|
| #Model_31_null | 4 | -180.65 | -173.17 | 94.327 | -188.65 | | | |
| #Model_31_glmm | 7 | -203.33 | -190.23 | 108.664 | -217.33 | 28.674 | 3 | **2.622e-06** |

**Pairwise contrast**

| contrast | estimate | SE | z.ratio | p.value |
|---|---|---|---|---|
| #C - NP | 0.08985 | 0.0163 | 5.518 | **<.0001** |
| #C - DP | 0.01086 | 0.0163 | 0.667 | 0.9095 |

| | | | | |
|---|---|---|---|---|
| #C - LP | 0.00541 | 0.0163 | 0.332 | 0.9873 |
| #NP - DP | -0.0789 | 0.0163 | -4.852 | **<.0001** |
| #NP - LP | -0.0844 | 0.0163 | -5.185 | **<.0001** |
| #DP - LP | -0.0054 | 0.0163 | -0.335 | 0.9871 |

Model_T2_glmm

| | npar | AIC | BIC | logLik | deviance | Chisq | Df | Pr(>Chisq) |
|---|---|---|---|---|---|---|---|---|
| #Model_T2_null | 4 | -16.731 | -9.2461 | 12.365 | -24.731 | | | |
| #Model_T2_glmm | 7 | -20.200 | -7.1016 | 17.100 | -34.200 | 9.4692 | 3 | **0.02366** |

**Pairwise contrast**

| contrast | estimate | SE | z.ratio | p.value |
|---|---|---|---|---|
| #C - NP | 1.372 | 0.0809 | 2.898 | **0.0197** |
| #C - DP | 1.036 | 0.0809 | 0.324 | 0.9882 |
| #C - LP | 1.018 | 0.0809 | 0.163 | 0.9984 |
| #NP - DP | 0.755 | 0.0809 | -2.575 | **0.04012** |
| #NP - LP | 0.742 | 0.0809 | -2.737 | **0.0315** |
| #DP - LP | 0.983 | 0.0809 | -0.161 | 0.9985 |

**Supplementary Table 5.** 16S rRNA beta diversity PERMANOVA and pairwise treatment comparisons calculated on Unweighted and Weighted dissimilarity matrices.

| **Overall** | **df** | **R2** | **F** | **Pr(>F)** |
|---|---|---|---|---|
| **Unweighted Unifrac** | 6 | 0.448 | 1.157 | 0.044 |
| **Weighted Unifrac** | 6 | 0.034 | 2.14 | 0.018 |

| **PAIRWISE** | **T0** | | | | **T1** | | | | **T2** | | | |
|---|---|---|---|---|---|---|---|---|---|---|---|---|
| **Unweighted Unifrac** | **df** | **R2** | **F** | **p-adj.** | **df** | **R2** | **F** | **p-adj.** | **df** | **R2** | **F** | **p-adj.** |
| #C - NP | 1 | 0.044 | 0.880 | 0.93 | 1 | 0.036 | 0.785 | 0.931 | 1 | 0.052 | 0.999 | 0.808 |
| #C - DP | 1 | 0.041 | 0.806 | 0.93 | 1 | 0.083 | 1.920 | **0.002** | 1 | 0.045 | 0.905 | 0.969 |
| #C - LP | 1 | 0.043 | 0.919 | 0.93 | 1 | 0.081 | 2.310 | **0.002** | 1 | 0.049 | 1.030 | 0.808 |
| #NP - DP | 1 | 0.051 | 0.972 | 0.93 | 1 | 0.103 | 1.690 | **0.002** | 1 | 0.041 | 0.820 | 0.969 |
| #NP - LP | 1 | 0.049 | 0.978 | 0.93 | 1 | 0.090 | 1.690 | **0.004** | 1 | 0.049 | 1.040 | 0.808 |
| #DP - LP | 1 | 0.056 | 1.130 | 0.93 | 1 | 0.059 | 1.080 | 0.254 | 1 | 0.041 | 0.895 | 0.969 |
| **Unweighted Unifrac** | **df** | **R2** | **F** | **p-adj.** | **df** | **R2** | **F** | **p-adj.** | **df** | **R2** | **F** | **p-adj.** |
| #C - NP | 1 | 0.048 | 0.962 | 0.588 | 1 | 0.016 | 0.353 | 0.923 | 1 | 0.045 | 0.851 | 0.848 |
| #C - DP | 1 | 0.046 | 0.928 | 0.588 | 1 | 0.182 | 4.690 | **0.012** | 1 | 0.045 | 0.901 | 0.848 |
| #C - LP | 1 | 0.024 | 0.502 | 0.936 | 1 | 0.248 | 5.930 | **0.015** | 1 | 0.052 | 1.110 | 0.848 |
| #NP - DP | 1 | 0.053 | 1.020 | 0.588 | 1 | 0.183 | 4.470 | **0.015** | 1 | 0.036 | 0.713 | 0.848 |
| #NP - LP | 1 | 0.024 | 0.468 | 0.936 | 1 | 0.249 | 5.650 | **0.006** | 1 | 0.045 | 0.953 | 0.848 |
| #DP - LP | 1 | 0.048 | 0.974 | 0.588 | 1 | 0.059 | 1.070 | 0.468 | 1 | 0.037 | 0.800 | 0.848 |

**Supplementary Table 6.** 16S rRNA alpha diversity metrics expressed as averages (± standard error) per treatment per timepoint of the study.

| Sampling time | Treatment | ASVs number | Chao1 | Shannon (H') |
|---|---|---|---|---|
| T0 | C | 526.83 (±13.11) | 849.51 (±21.54) | 3.95 (±0.07) |
| | NP | 480.83 (±32.29) | 775.07 (±46.74) | 3.97 (±0.18) |
| | LP | 516.50 (±33.66) | 818.46 (±46.15) | 3.89 (±0.09) |
| | DP | 514.92 (±17.32) | 846.72 (±24.52) | 3.79 (±0.01) |
| T1 | C | 475.08 (±37.71) | 818.01 (±92.41) | 3.42 (±0.18) |
| | NP | 468.92 (±27.87) | 745.43 (±40.24) | 3.47 (±0.19) |
| | LP | 595.67 (±50.22) | 841.87 (±50.59) | 4.33(±0.17) |
| | DP | 541.17 (±46.98) | 804.73 (±48.42) | 4.02 (±0.17) |
| T2 | C | 1602.50 (±365.98) | 2161.64 (±478.11) | 6.04 (±0.17) |
| | NP | 1909.50 (±420.25) | 2702.65 (±787.13) | 6.26 (±0.18) |
| | LP | 1688.83 (±294.12) | 2143.87 (±356.66) | 6.26 (±1.84) |
| | DP | 1711.25 (±345.69) | 2579.70 (±918.42) | 6.19 (±0.11) |

**Supplementary Table 7.** Statistical comparisons between treatments for alpha diversity metrics (ASVs number, Shannon index, Chao1 index) calculated by implementing the Dunn test for multiple comparisons.

| | Kruskal-Wallis | ASVs number | | |
|---|---|---|---|---|
| Overall | T0 | chi-squared = 4.88 | df = 3 | p = 0.18 |
| | T1 | chi-squared = 1.80 | df = 3 | p = 0.62 |
| | T2 | chi-squared = 2.72 | df = 3 | p = 0.44 |
| | Dunn's Test | T0 | T1 | T2 |
| Pairwise contrasts | C-NP | p = 0.17 | p = 0.22 | p = 0.07 |
| | C-LP | p = 0.08 | p = 0.39 | p = 0.09 |
| | C-DP | p = 0.20 | p = 0.29 | p = 0.09 |
| | NP-LP | p = 0.23 | p = 0.17 | p = 0.44 |
| | NP-DP | p = 0.10 | p = 0.10 | p = 0.43 |
| | LP-DP | p = 0.29 | p = 0.41 | p = 0.49 |
| | **Kruskal-Wallis** | **Shannon (H')** | | |
| Overall | T0 | chi-squared = 1.76 | df = 3 | p = 0.62 |
| | T1 | chi-squared = 13.50 | df = 3 | **p < 0.01** |
| | T2 | chi-squared = 2.45 | df = 3 | p = 0.48 |
| | Dunn's Test | T0 | T1 | T2 |
| Pairwise contrasts | C-NP | p = 0.27 | p = 0.44 | p = 0.07 |
| | C-LP | p = 0.39 | **p < 0.01** | p = 0.32 |
| | C-DP | p = 0.10 | **p < 0.01** | p = 0.42 |
| | NP-LP | p = 0.37 | **p < 0.01** | p = 0.15 |
| | NP-DP | p = 0.26 | **p < 0.01** | p = 0.10 |
| | LP-DP | p = 0.16 | p = 0.35 | p = 0.38 |
| | **Kruskal-Wallis** | **Chao1** | | |
| Overall | T0 | chi-squared = 1.39 | df = 3 | p = 0.71 |
| | T1 | chi-squared = 4.07 | df = 3 | p = 0.25 |
| | T2 | chi-squared = 1.37 | df = 3 | p = 0.71 |
| | Dunn's Test | T0 | T1 | T2 |
| Pairwise contrasts | C-NP | p = 0.12 | p = 0.46 | p = 0.15 |
| | C-LP | p = 0.26 | p = 0.10 | p = 0.17 |
| | C-DP | p = 0.23 | p = 0.07 | p = 0.32 |
| | NP-LP | p = 0.29 | p = 0.10 | p = 0.44 |
| | NP-DP | p = 0.29 | p = 0.05 | p = 0.27 |
| | LP-DP | p = 0.45 | p = 0.43 | p = 0.31 |

**Supplementary Table 8.** Average relative abundances (± standard error) of bacterial families per treatment per timepoint of the study (Excel file: "Families_Rel_abundances.xlsx").
Available upon publication at 10.5281/zenodo.17279049.

**Supplementary Table 9.** Complete list of the differentially abundant ASVs resulting from ANCOM-BC2 analysis (including "Unclassified" ASVs) at each timepoint of the experiment Excel file: "Ancombc2_results.xlsx").
Available upon publication at 10.5281/zenodo.17279049.

**Supplementary Table 10.** Host transcriptomic profiles PERMANOVA results.

| | **df** | **R2** | **F** | **Pr(>F)** |
|---|---|---|---|---|
| **Overall** | 3 | 0.047 | 0.849 | **0.012** |
| **PAIRWISE** | **T0-T1-T2** | | | |
| | **df** | **R2** | **F** | **p-adj.** |
| #C - NP | 2 | 0.191 | 2.501 | **0.0001** |
| #C - DP | 2 | 0.186 | 2.353 | **0.001** |
| #C - LP | 2 | 0.198 | 2.548 | **0.001** |
| #NP - DP | 2 | 0.22 | 2.98 | **8.00E-05** |
| #NP - LP | 2 | 0.223 | 3.016 | **5.00E-05** |
| #DP - LP | 2 | 0.229 | 3.053 | **0.0002** |

| **PAIRWISE** | **T0** | | | | **T1** | | | | **T2** | | | |
|---|---|---|---|---|---|---|---|---|---|---|---|---|
| | **df** | **R2** | **F** | **p-adj.** | **df** | **R2** | **F** | **p-adj.** | **df** | **R2** | **F** | **p-adj.** |
| #C - NP | 1 | 0.046 | 0.293 | 0.942 | 1 | 0.116 | 0.791 | 0.685 | 1 | 0.199 | 1.497 | 0.171 |
| #C - DP | 1 | 0.171 | 0.548 | 0.714 | 1 | 0.079 | 0.517 | 0.685 | 1 | 0.134 | 0.929 | 0.542 |
| #C - LP | 1 | 0.058 | 0.375 | 0.8 | 1 | 0.106 | 0.716 | 0.657 | 1 | 0.137 | 0.959 | 0.342 |
| #NP - DP | 1 | 0.062 | 0.4 | 1 | 1 | 0.125 | 0.857 | 0.571 | 1 | 0.124 | 0.849 | 0.685 |
| #NP - LP | 1 | 0.047 | 0.296 | 0.971 | 1 | 0.12 | 0.819 | 0.6 | 1 | 0.172 | 1.252 | 0.214 |
| #DP - LP | 1 | 0.078 | 0.513 | 0.885 | 1 | 0.067 | 0.436 | 0.857 | 1 | 0.087 | 0.57 | 0.942 |

**Supplementary Table 11.** Complete list of the differentially expressed genes (DEGs) resulting from DESeq2 differential gene expression analysis (Excel file: "DEGs.xlsx").
Available upon publication at 10.5281/zenodo.17279049.

**Supplementary Table 12.** Complete list of the GO terms resulting from the gene ontology enrichment analysis (Excel file: "GO. xlsx").
Available upon publication at 10.5281/zenodo.17279049.
